# The Immune Response Against Cancer is Modulated by Stromal Cell Fibronectin

**DOI:** 10.1101/2025.05.14.653424

**Authors:** Alexander Lubosch, Lauren Pitt, Caren Zoeller, Franziska Wirth, Tarik Exner, Barbara Steigenberger, Jutta Schroeder-Braunstein, Guido Wabnitz, Inaam A. Nakchbandi

## Abstract

Cancer-associated fibroblasts remain poorly understood, with some of them originating from the bone marrow. We therefore took advantage of the diversity of bone marrow stromal cells to shed light on how fibroblasts modulate cancer growth.

In two murine cancer models, adding these fibroblasts to tumor cells resulted in smaller lesions. Suppression was enhanced by pretreatment with fibronectin, while genetic deletion of fibronectin in a small subpopulation of stromal cells expressing osterix/*sp7* restored growth. The suppressive stromal population showed two more characteristics: the absence of CD31/*pecam1* and CD105/*endoglin*. However, only a decrease in CD105/*ENDOGLIN* in melanoma patients translated in improved survival. Mechanistically, fibronectin or fibronectin fragments activate integrin α5β1 and TLR4 and increase chemokine production by stromal cells ultimately leading to enhanced recruitment and activity of Ly6G^+^ myeloid cells without T-cell involvement.

This work thus characterizes a beneficial interaction between stromal cells and neutrophils enhancing the immune response against early cancer.

**Highlights:**

- Stromal cells can be divided into two populations, a tumor supportive one with the potential to stimulate angiogenesis and an inhibitory one with the potential to enhance the immune response
- Stromal fibronectin expressing the transduction factor osterix/*sp7* produce fibronectin that acts on α5β1 integrin and/or TLR4 on neighboring fibroblasts to modulate Ly6G^+^ immune cells and suppress tumor growth

**One sentence summary:** Tumor inhibitory fibroblasts have three characteristics and through production of fibronectin act on neighboring cells inducing signaling cascades that ultimately lead to a stronger immune response against early cancer.

## Introduction

Tumor cells are the main actors in cancer, but other cells are involved in modulating lesion growth. These include immune, vascular and stromal cells. However, stromal cells in the tumor represent a heterogeneous population called cancer-associated fibroblasts (CAFs)^1^. There are no specific markers to define CAFs^2^, but the expansive variety of markers suggests that CAFs differ within a tumor and between different tumors, and that they may have different functions^1,3^.

Opposing effects on tumor growth have been attributed to CAFs. Cells expressing markers such as fibroblast-specific protein-1 (FSP1, also called S100A4) or fibroblast associated protein (FAP) were associated with increased growth^4,5^. In contrast, αSMA-expressing cells suppressed growth because their depletion enhanced the epithelial-mesenchymal transition of pancreatic cancer cells, leading to disease progression^6^. Another subpopulation capable of producing collagen I similarly inhibited cancer growth^7^.

An immunomodulatory ability was also ascribed to CAFs, suggesting that they may influence tumor growth by modifying immune cell number or behavior. The molecular mechanisms involved remain elusive, however. One possibility is that CAFs prejudice the immune cells to support cancer progression^8,9^. However, the opposite has also been reported. As an example, a subpopulation of stromal cells that did not express CD105 (endoglin, a cell surface protein) instructed dendritic cells and T lymphocytes to suppress tumor growth^10^.

Even though the role of the immune system in fighting cancer is complex, neutrophils have been gaining relevance. They can attack tumor cells through a variety of mechanisms such as the oxidative burst^11^. On the other hand, tumor cells can sometimes inhibit neutrophil migration and polarization relatively quickly leading to the inability of these cells to halt tumor progression^12^. CAFs were also shown to increase recruitment and survival of immune suppressive neutrophils^13^. Thus, there is experimental evidence for stromal cell-neutrophil interactions in tumor immunity, but the underlying mechanisms are still poorly understood.

There are several subtypes of fibroblasts. Mesenchymal stromal cells (MSCs) isolated from the bone marrow represent a heterogeneous group. These cells are particularly interesting, because some have both hematopoietic and fibroblastic properties^14,15^, a feature that could explain their potential to modify immune responses. We have shown that a subpopulation of stromal cells isolated from the bone marrow of healthy mice efficiently suppresses tumor growth^16^. Other groups showed that stromal cells can either inhibit or stimulate growth depending on the context^3^. Additionally, these cells seem to induce immune changes that lead to tumor growth enhancement^13^.

Our aim was to better understand the role of fibroblasts in tumor growth. We chose the bone marrow stromal cells as a source of heterogeneous primary fibroblasts and embarked on the search for a mechanism for their effect on tumors. We identified two subpopulations with opposite effects. Neutrophils were found to be crucial in our model, and their recruitment was dependent on the activation of stromal cells by fibronectin.

## Results

### Stromal cells from the bone marrow inhibit tumor growth

To better understand how fibroblasts modulate cancer growth, bone marrow stromal cells were used as a source of various fibroblastic cells with diverse functions. Non-hematopoietic cells isolated from the bone marrow were mixed with 10^6^ B16 melanoma cancer cells at a ratio of 1:0.1 (10^6^ B16 cancer cells:10^5^ bone marrow cells), 1:1 and 1:2 (Purity and viability of the cells are shown in Supplementary-Figure 1A-B). The mixture was injected subcutaneously and growth was evaluated (Figure 1A). Injection of stromal cells (SCs) at a ratio of 1:0.1 inhibited growth, while an increase in stromal cells in relation to tumor cells (1:2) resulted in growth similar to that in the absence of fibroblasts (Figure-1B and Supplementary-Figure 1C). Since we aimed to understand how suppression is mediated, we performed all subsequent experiments using the inhibitory ratio of 1:0.1.

**Figure 1.**
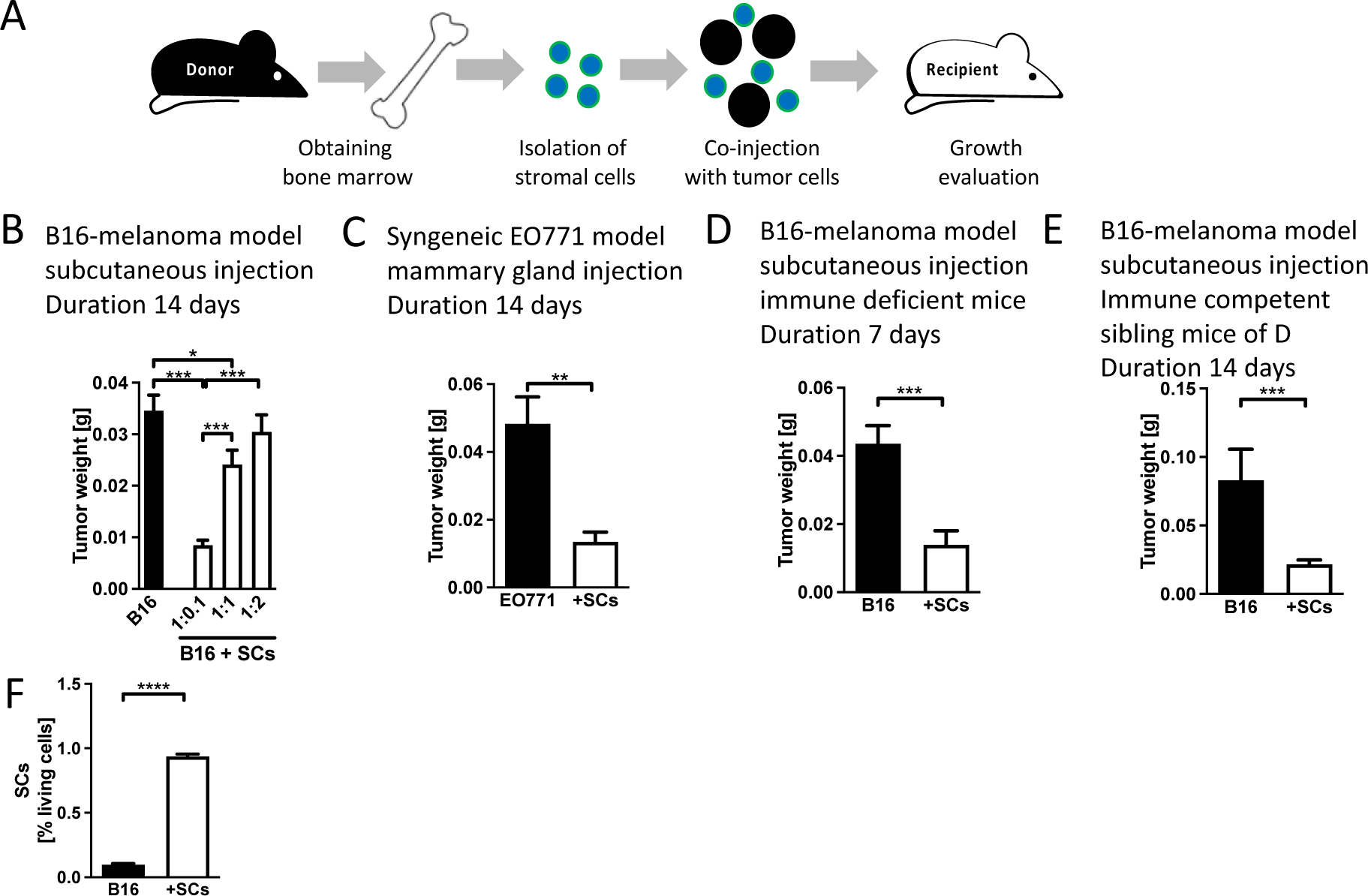
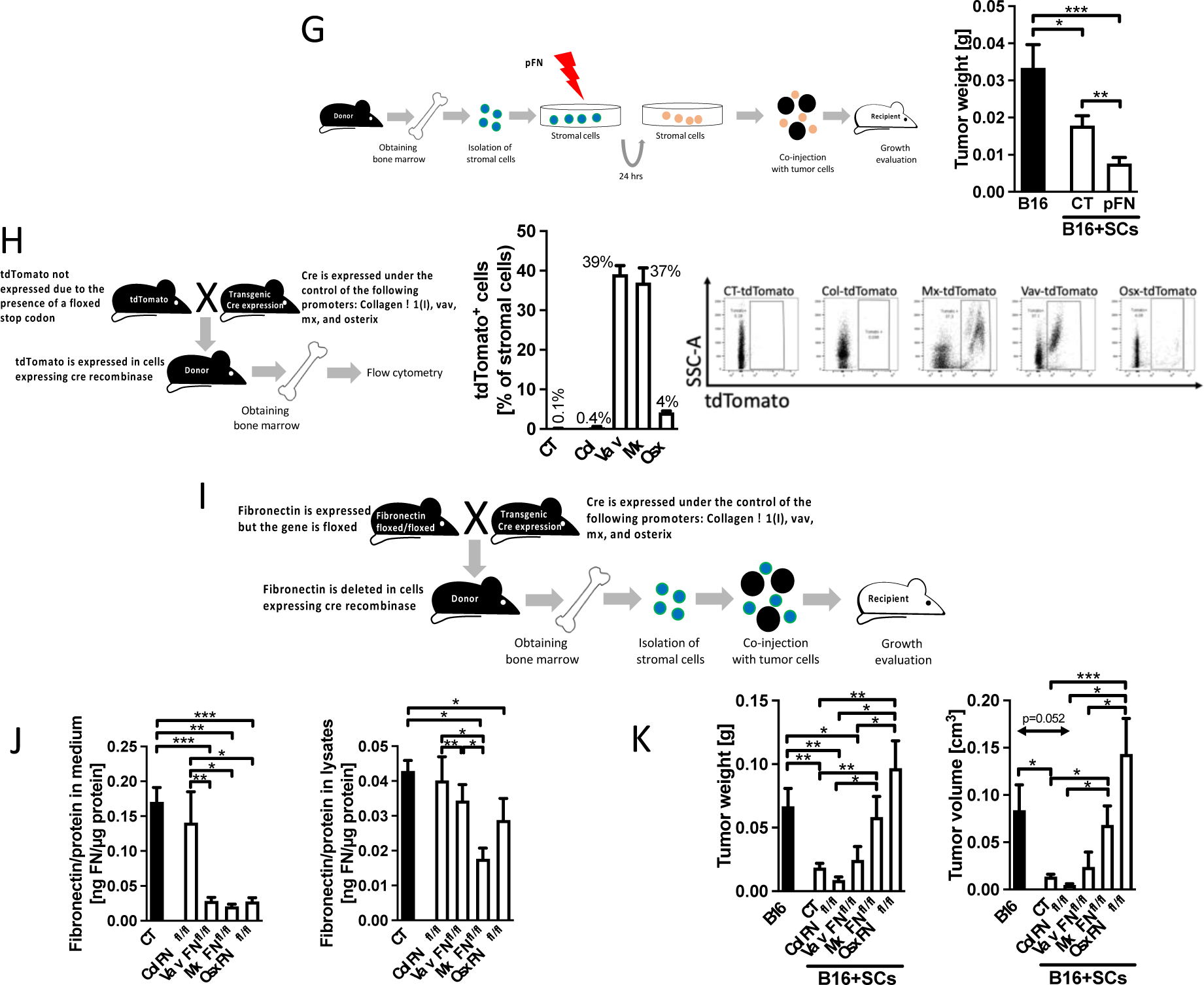
Bone marrow stromal cells suppress tumor growth G-K. Fibronectin mediates suppression of growth. A. Schematic of the experiment. Bone marrow cells were isolated, depleted of immune cells, and mixed in different ratios with tumor cells before injecting the mixture subcutaneously. Growth was evaluated after 7-14 days. B. Injection of stromal cells with B16 melanoma tumor cells at a ratio of 1:0.1 (Tumor cells:stromal cells) results in smaller tumors. The ratio of 1:1 still led to a small decrease in tumor weight, and the 1:2 ratio failed to inhibit cancer growth. N=12/12/12/12, *p<0.05, ***p<0.001. C. Using syngeneic breast tumor cells (EO771) at a ratio of 1:0.1 injected orthotopically in the mammary gland confirms diminished growth. N=10/8, **p<0.01, ***p<0.001. D. Tumor suppression in the presence of stromal cells is maintained using B16 melanoma cells in animals lacking T cells. N=8/8, **p<0.01, ***p<0.001. E. A different genetic background of the mice did not prevent inhibition by stromal cells. N=9/15, **p<0.01. F. Stromal cells were labeled before mixing with B16 tumor cells and injection subcutaneously. Three days after injection only 1% of the cells in the tumor were labeled suggesting that few cells are sufficient for the inhibitory effect. N=4/5, *p<0.05. G. Schematic of pretreatment of stromal cells (SCs) before injecting them. After isolation of bone marrow cells and depletion of immune cells, stromal cells were cultured for 24 hours in the presence of plasma fibronectin at a concentration of 160 μg/ml. Afterwards, these cells were mixed with tumor cells and injected into mice. Pretreatment of stromal cells with fibronectin inhibits tumor growth more than untreated stromal cells. N=6/7/9, *p<0.05, **p<0.01, ***p<0.001. H. Schematic for detection of Cre expression using tdTomato transgenic mice. Mice carrying the gene for cre-recombinase under the control of the various promoters were mated with animals homozygous for tdTomato. Half the pups were controls and half carried the promoter attached to cre in addition to the gene for tdTomato. The tdTomato gene is preceded by a stop codon that is floxed. Expression of cre-recombinase in cells that also contain the transgene tdTomato leads to removal of the stop codon and production of fluorescent tdTomato. Flow cytometry confirms expression of cre-recombinase in bone marrow stromal cells except in the case of Colα1(I). N=5/4/3/4/4. I. Schematic of conditional deletion of fibronectin in transgenic mice using mice homozygous for floxed fibronectin and mice carrying the cre-recombinase gene under the control of the promoters used. Matings over two generations allow deletion of fibronectin in stromal cells and use of these cells in tumor growth experiments. J. Fibronectin deletion in various bone marrow stromal cells. Vav, mx and osx showed a strong suppression of fibronectin in the conditioned media after 24 hours in culture. In cell lysates, osx led to decreased fibronectin despite its expression in only 4% of bone marrow cells. Total bone marrow stromal cells after depletion of immune cells were evaluated. N=41/9/7/14/11, *p<0.05, **p<0.01, ***p<0.001. K. Freshly isolated stromal cells from the bone marrow of transgenic mice showed suppression of growth except in the case of stromal cells from mice in which fibronectin was deleted in mx or in osterix cells. N=16/16/8/11/10/13, *p<0.05, **p<0.01, ***p<0.001. Comparisons were performed using ANOVA, and if significant, were followed by t-tests or t-tests only as appropriate.

To determine whether this ratio is able to suppress tumor growth in other models, we injected orthotopic syngeneic EO771 cells, a murine breast cancer cell line, into the mammary gland and confirmed suppressed growth (Figure 1C and Supplementary-Figure 1D).

Since T cells to mediate inhibitory effects of stromal cells on tumor growth^13^, we evaluated the percentage of immune cells and T cells and found no difference. This suggests that a change in T-cell numbers is not responsible for diminished growth in the presence of stromal cells (Supplementary-Figure 1E-F) ^10^. To study this further, mice homozygous for the *foxn1nu* (nu/nu) mutation and hence unable to mount T cell-mediated immune responses were used^17^. Repeating the experiments using B16 melanoma cells confirmed suppression of growth despite the lack of mature T cells (Figure 1D and Supplementary-Figure 1G). Since these animals had a mixed genetic background, the experiments were performed in parallel in immune competent littermate controls which confirmed that the decrease in growth is independent of the background (Figure 1E and Supplementary-Figure 1H).

Next, we evaluated whether stromal cells could still be detected in the developing tumors after 3 days. Surprisingly, less than 1% of pre-labeled stromal cells were found in the tumors by flow cytometry (Figure 1F and Supplementary-Figure 1I). This suggests that the inhibitory effect of stromal cells is initiated at very early stages of tumor development.

To exclude a role of contaminating immune cells (<2%, Supplementary-Figure 1A), CD45^+^ immune cells were mixed with melanoma B16 cells at a ratio of 1:0.1 (Supplementary-Figure 2A). Similarly, freshly sorted CAFs from a human breast cancer model in mice were applied at the same ratio (Supplementary-Figure 2B) (The sorting method is shown in Supplementary-Figure 2C). Under both conditions, cancer growth was not suppressed.

In summary, non-hematopoietic stromal bone marrow cells suppress tumor growth in two different tumor models. This suppression does not require the presence of functional T cells.

### Fibronectin in stromal cells is required for suppression of growth

#### Fibronectin addition

Fibronectin supports tumor growth as shown using several models^18–20^. It is also produced by a variety of stromal cells^21^. We therefore sought to determine whether pretreating bone marrow stromal cells with fibronectin for 24 hours prior to mixing with cancer cells and injecting into mice would enhance tumor growth. Interestingly, the opposite occurred, showing that treatment of isolated bone marrow stromal cells with plasma fibronectin (the isoform lacking the extra domains A and B) suppressed growth even more (Figure 1G).

#### Fibronectin deletion

We therefore asked whether deleting fibronectin in various bone marrow stromal cells would prevent their suppressive effects. The cre/loxP system allows use of promoters attached to cre-recombinase to delete fibronectin in animals that are homozygous for the floxed fibronectin gene. The following promoters known to be expressed in bone marrow stromal cells were evaluated: vav (detected in hematopoietic and non-hematopoietic cells); mx (which affects many different cell types), and osterix/sp7 (Osx) (expressed in early osteoblasts and chondrocyte progenitors)^22–24^. Collagen α1(I), which characterizes differentiating osteoblasts, was also used^25^. Since only stromal cells were employed (after depletion of CD45^+^ hematopoietic cells), any effect of these promoters on hematopoietic cells is not relevant in our model.

To determine the activity of the promoters in stromal cells, we took advantage of tomato reporter mice (tdTomato)^22^. Cells in which Cre is expressed will remove a floxed stop codon preceding the tdTomato gene, produce the protein tdTomato and, consequently, allow detection of Cre expressing cells by flow cytometry (Figure 1H). Flushed bone marrow stromal cells expressed the evaluated promoters except for Colα1(I), which is active in osteoblasts on the bone surface and therefore not detected in bone marrow cell isolates. Indeed, only 4% of stromal cells expressed osterix, which is found in chondrocytic and osteoblastic precursors (figure 1H)^23,26^. Deletion of total fibronectin in bone marrow stromal cells was successful to various degrees, except when using the Colα1(I) promoter. Stromal cells from the Colα1(I) mice can therefore serve as Cre controls (Figure 1I-J). Notably, fibronectin in the cell lysates was only diminished in mx mice (Mx-FN*^fl/fl^*: Mx-cre_fibronectin*^floxed/floxed^*) and osterix mice (Osx-FN*^fl/fl^*: Osterix-cre_fibronectin*^floxed/floxed^*) (Figure 1J, last two bars of right graph).

Stromal cells isolated from the various transgenic mouse models of fibronectin deletion were used. A decrease in growth was seen except when fibronectin was deleted using the mx or osterix promoters, in Mx-FN*^fl/fl^* and Osx-FN*^fl/fl^* mice (Figure 1K), which were the only mouse lines with confirmed deletion of fibronectin in cell lysates (Figure 1J, right graph). It is possible that the osterix subpopulation is included in the larger mx subpopulation, because the mx promoter also targets osteogenic cells^22^. Because mx, like vav, is expressed in many stromal cells, we focused on osterix-expressing cells found in only 4% of stromal cells (Figure 1H).

In summary, fibronectin produced by a subpopulation that expresses the transcription factor osterix/sp7 is required for tumor suppression.

### Evaluating integrins as possible mediators of stromal suppression

Several members of the integrin family bind to fibronectin, and some of them contain the β1 subunit^27^. We therefore examined whether deletion of β1 integrin in bone marrow cells affected growth suppression by stromal cells. By flow cytometry, deletion of β1 integrin was only measurable in Vav-β1 (Vav-cre_β1*^floxed/floxed^*) and Mx-β1*^fl/fl^* (Mx-cre_β1*^floxed/floxed^*) conditional knockout mice (Figure 2A). Since Osx is expressed only in a small percentage of bone marrow cells, we sorted osterix-expressing cells by taking advantage of the tomato reporter gene after introducing it in Osx-β1 *^fl/fl^* mice (Genotype: Osx-cre_tdTomato*^fl/+^*_β1*^floxed/floxed^*). This confirmed diminished β1 integrin (but not complete deletion) by flow cytometry and western blotting (Figure 2B-D). Mixing bone marrow stromal cells from Osx-β1 *^fl/fl^* with tumor cells resulted in smaller lesions, similar to control cells. Interestingly, only deletion of β1 in Mx-β1 *^fl/fl^* mice led to larger tumors than all other models, but growth was not fully restored (Figure 2E), unlike fibronectin deletion in osterix or mx mice (Figure 1K). This could be either due to the absence of a role of a β1 integrin in the Osx cells or due to the incomplete deletion of β1 integrin in this population compared to Mx-β1 *^fl/fl^*. In summary, β1 may play a role in mediating suppression of growth.

**Figure 2.**
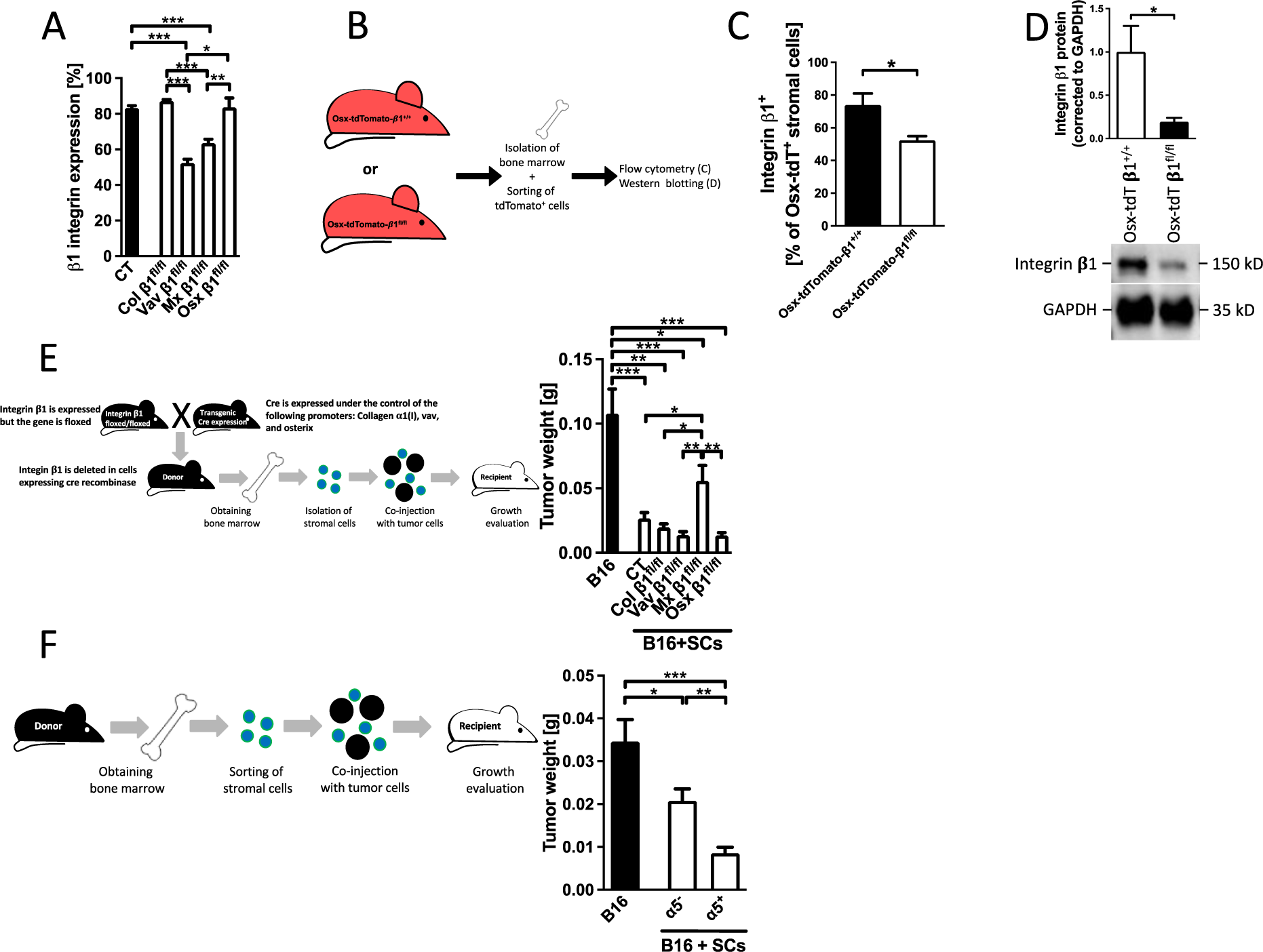
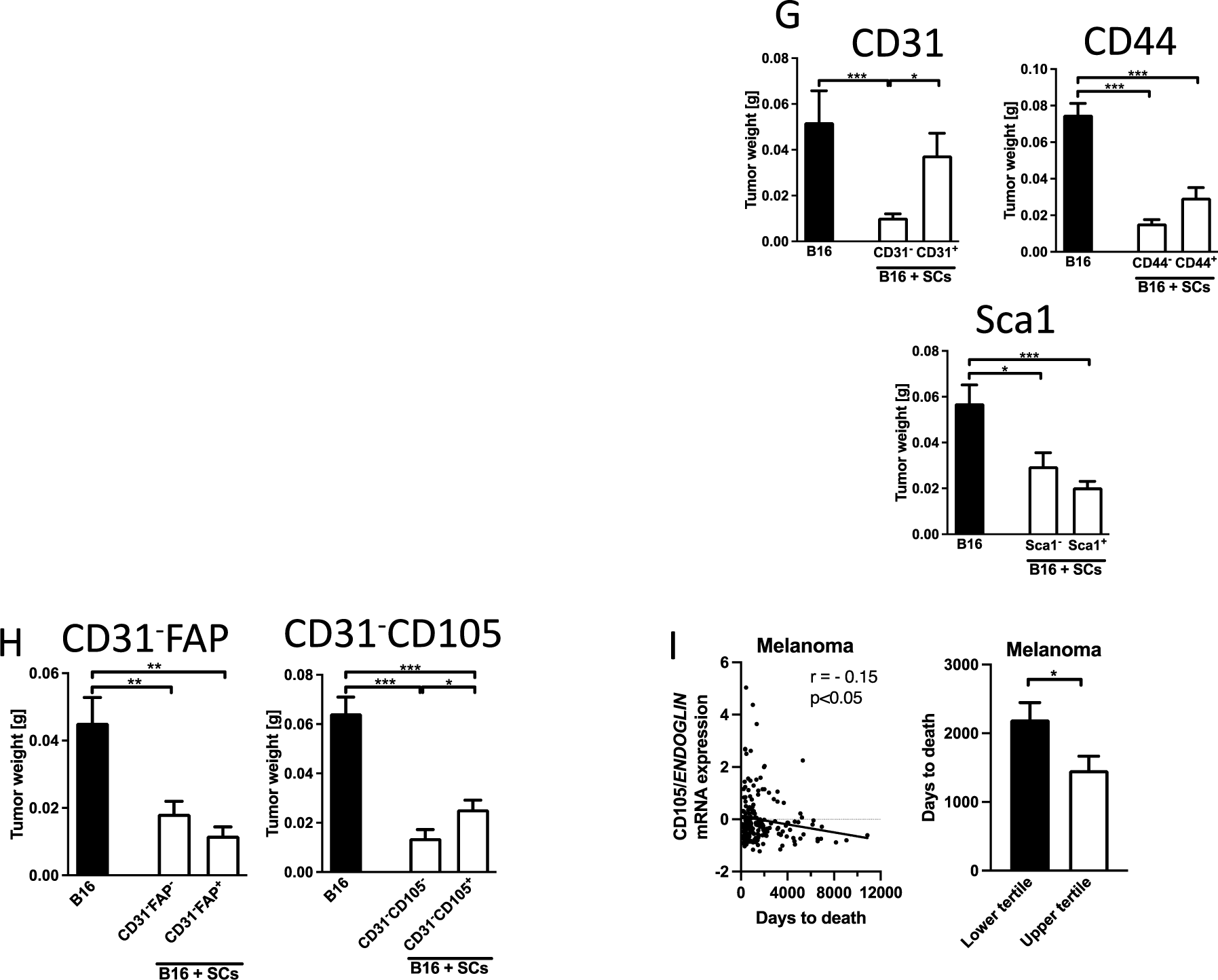
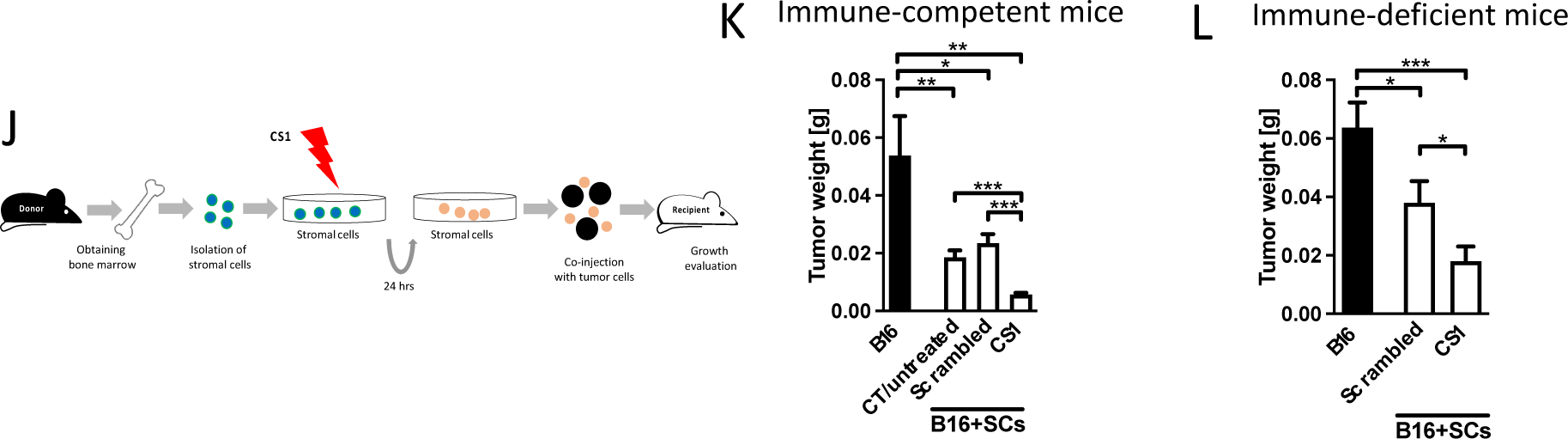
The role of Integrin α5β1 in suppression of growth G-H. Sorted stromal cells differ in their inhibitory effects on tumor growth. J-L. CS1 enhances inhibition by stromal cells. <u>**A-E. β1 subunit**</u>: A. Expression of integrin subunit β1 on the bone marrow cells of transgenic mouse models in which β1 was depleted using the promoters colα1(I), vav, mx and osterix. Bone marrow was isolated and stained. Cells that expressed neither CD45 nor Ter119, but stained for β1 integrin are shown. The percentages thus represent the expression of β1 subunit in stromal cells (SCs). N=13/10/2/7/3, *p<0.05, ***p<0.001. B. Schematic showing the two genotypes evaluated for confirmation of depletion of β1 integrin. Mice were mated to generate β1 conditional knockout mice that express tdTomato whenever the promoter was expressed. If the cells contain another floxed gene (in this case the integrin subunit β1), the gene will also be deleted. Consequently, tdTomato-labeles cells should also lack β1. The control mice had the osterix promoter attached to cre, a copy of the tdTomato/^+^ gene and wildtype β1 integrin genes. The bone marrow was isolated, stained, sorted, and evaluated. C. Stromal cells from Osx-tdTomato^+^-β1^fl/fl^ show a decrease but not complete deletion of β1 compared to Osx-tdTomato/^+^-β1^+/+^. N=2/6, *p<0.05. D. Sorted bone marrow cells from Osx-tdTomato/^+^-β1^fl/fl^ show a decrease in protein expression of β1 compared to controls (Osx-tdTomato/^+^-β1^+/+^). N=2/4, *p<0.05. E. Schematic showing deletion of β1 in transgenic mice using mice homozygous for floxed integrin subunit β1 and mice carrying the cre-recombinase gene under the control of different promoters. Matings over two generations allow deletion of β1 subunit in stromal cells and use of these cells in tumor growth experiments. Stromal cells from transgenic mice showed suppression of growth in all models except when using the mx promoter, where growth increased but was not fully restored. N=17/16/9/12/15/11, *p<0.05, **p<0.01, ***p<0.001. **<u>F. α5 subunit:</u>** F. Schematic of experiment on the role of integrin α5 subunit. Stromal cells (SCs) from wildtype mice were isolated, depleted of immune cells, stained and sorted for α5 integrin subunit, and mixed with B16 tumor cells at a ratio of 1:0.1 (B16/SCs) before injecting them into mice subcutaneously. Growth was evaluated after 14 days. Inhibition of tumor growth by stromal cells was more pronounced in cells that express α5 and hence are able to respond to fibronectin, but the α5^-^ cells also suppressed growth. N=19/19/18, *p<0.05, **p<0.01, ***p<0.001. Inhibition of tumor growth by stromal cells was lost in CD31^+^ cells, but the other evaluated subpopulations suppressed growth to various degrees (G). The addition of FAP or CD105 to CD31^-^ inhibitory cells revealed enhanced growth suppression in the absence of CD105 (H). The evaluated population is highlighted in the title of each graph. CD31: N=4/10/9; Sca1: N=5/11/10; CD44: N=6/9/9; CD31-FAP: N=6/11/6 ; CD31-CD105: N=6/12/12, *p<0.05, **p<0.01, ***p<0.001. **I. Human study.** Gene expression analysis from primary skin melanoma lesions from 193 patients without prior malignancy, prior treatment, or synchronous malignancy shows a negative correlation between CD105/ENDOGLIN gene expression and the number of days to death. *p<0.05. The patients in the lower tertile for gene expression have longer survival than those in the upper tertile. N=193. Pearson’s correlation and t-test were used for comparison. *p<0.05. J. Schematic of experiment. Isolated bone marrow cells were depleted from immune cells. Stromal cells (SCs) were pretreated for 24 hours in vitro, mixed with tumor cells and injected into mice. K. Pretreatment of stromal cells with CS1 suppressed tumor growth more so than untreated stromal cells or cells pretreated with a scrambled control peptide. The scrambled peptide and CS1 were used at a concentration of 20 μg/ml. N=14/15/12/15, *p<0.05, **p<0.01, ***p<0.001. L. CS1 pretreatment inhibits cancer growth despite the absence of mature T cells. N=12/10/11, *p<0.05, **p<0.01, ***p<0.001. The experiment was performed as shown in J, but in mice that are homozygous for a foxn1 mutation (foxn1^nu/nu^). They therefore lack a thymus and functional T cells. Comparisons were performed using ANOVA followed by t-tests.

Integrin α5β1 represents the classical fibronectin receptor^28^. In order to determine whether the presence of α5 integrin on stromal cells leads to diminished cancer growth, we sorted α5 expressing stromal cells (α5^+^) and cells not expressing this receptor subunit (α5^-^), mixed these with cancer cells, and injected them. Inhibition of growth was more pronounced when using α5^+^ cells that express α5, albeit also present when using α5^-^ cells, suggesting a limited role of α5 in mediating growth suppression (Figure 2F).

Thus, a small subpopulation of approximately 4% of the bone marrow stromal cells (expressing osterix) produces fibronectin, which is able to suppress tumor growth, but the inhibitory action of fibronectin on stromal cells is mediated only in part by α5β1 integrin.

### Characterization of osterix cells

To further characterize the osterix subpopulation, we examined stromal cells from tdTomato reporter mice that expressed osterix-cre (Osx-cre_tdTomato*^floxed/+^*; mice were generated as shown in Figure 1H).

These cells were first sorted, and their proteins were compared to stromal cells not expressing tdTomato. Proteomic analysis revealed that osterix-Cre expressing cells have characteristics of embryonic fibroblastic cells. Of the 30 molecules that differed highly significantly (p<0.01) and showed a 4-fold change between control and osterix-expressing cells, 11 were associated with connective tissue. This is not surprising since osterix cells are progenitors of osteoblasts and chondrocytes. Nine molecules were associated with nucleotide metabolism, consistent with DNA and RNA modulation. Six molecules were associated with protein folding and 6 with cellular response to stress. The volcano plot highlighted three molecules, all of which were related to proliferation (Ranbp1, HPRT1, and angptl3) (Supplementary-Figure 3 shows a heat map and a volcano plot; a list of the differentially expressed molecules is presented in Supplementary-Table 1). This suggests that osterix-expressing stromal cells respond differently to proliferative cues than other stromal cells.

We next stained stromal cells from Osx-tdTomato mice with various stromal cell markers. CD31, CD44, and CD105, and CD140b are expressed on vascular cells^16^; and fibroblast activating protein-α (FAP) is expressed on bone marrow stromal cells and on CAFs ^14^. Finally, Sca-1 defines a population with stem cell characteristics^16^. Comparing stromal cells that express osterix (Osx^+^) (from Osx-cre_tdTomato*^floxed/+^*) with those that did not (Osx^-^) showed that a higher percentage of osterix cells did not express CD31/PECAM1 (CD31^-^) and expressed the following markers: CD44^+^, CD105^+^, CD140b^+^, CD146^+^, LepR^+^, FAP^+^ and Sca1^+^. However, the expression of CD140b^+^, CD146^+^ and LepR^+^ on Osx^-^ stromal cells (representing 96% of the stromal cells) was less than 2%, making *in vivo* evaluations challenging. These three molecules were therefore excluded from further studies (Table 1).

**Table 1.** Surface marker expression on stromal cells in cells that do not express osterix (Osx^-^, 96% of total stromal cells) and in cells that express osterix (Osx^+^, 4%). Stromal cells were isolated from the bone marrow of Osx-tdTomato reporter mice (Genotype: Osx-cre_tdTomato^floxed/+^, method explained in Figure 1H and its legend). Using flow cytometry, osterix expression was detected by tdTomato label and the surface markers were evaluated after staining. N= 4. PDPN: podoplanin.

|  | CD31<sup>-</sup> | CD31<sup>+</sup> | CD44<sup>-</sup> | CD44<sup>+</sup> | CD90<sup>-</sup> | CD90<sup>+</sup> |
| --- | --- | --- | --- | --- | --- | --- |
| **Osx<sup>-</sup>** | 22.2 ± 2.9 | 74.8 ± 3.4 | 72.7 ± 9 | 27.1 ± 9 | 95.0 ± 1.8 | 4 ± 1.5 |
| **Osx<sup>+</sup>** | **34.2 ± 5.3** | 63.7 ± 5.5 | 69.3 ± 7.5 | **30.2 ± 7.4** | 93.8 ± 2.4 | 4.9 ± 2 |
|  | CD105<sup>-</sup> | CD105<sup>+</sup> | CD140a<sup>-</sup> | CD140a<sup>+</sup> | CD140b<sup>-</sup> | CD140b<sup>+</sup> |
| **Osx<sup>-</sup>** | 92.2 ± 1.2 | 7.1 ± 1 | 93.6 ± 3.9 | 6 ± 3.766 | 97.9 ± 0.6 | 1.6 ± 0.4 |
| **Osx<sup>+</sup>** | 78.9 ± 2.2 | **20.4 ± 2.1** | 93.3 ± 3.9 | 6.4 ± 3.8 | 86.2 ± 2.9 | **12.3 ± 2.4** |
|  | CD146<sup>-</sup> | CD146<sup>+</sup> | LepR<sup>-</sup> | LepR<sup>+</sup> | FAP<sup>-</sup> | FAP<sup>+</sup> |
| **Osx<sup>-</sup>** | 98.2 ± 1 | 1.6 ± 1 | 99.7 ± 0.1 | 0.3 ± 0.1 | 95.3 ± 2 | 4.3 ± 1.7 |
| **Osx<sup>+</sup>** | 95.8 ± 0.6 | **3.1 ± 0.6** | 90.1 ± 3.3 | **8.4 ± 2.9** | 78.7 ± 4.9 | **19 ± 4.5** |
|  | PDPN<sup>-</sup> | PDPN<sup>+</sup> | Sca1<sup>-</sup> | Sca1<sup>+</sup> |  |  |
| **Osx<sup>-</sup>** | 95.3 ± 2 | 4.3 ± 1.7 | 93.3 ± 1.7 | 6.3 ± 1.5 |  |  |
| **Osx<sup>+</sup>** | 94.2 ± 1.3 | 5.4 ± 1.2 | 87.8 ± 2.8 | **12.7 ± 2.8** |  |  |

### Characterization of the inhibitory population

In order to determine which stromal subpopulation mediates growth inhibition, we took advantage of the differential expression of the selected markers. Of the five subpopulations that were increased in Osx-expressing cells, three, namely CD31^-^, Sca1^+^, and CD44^+^ were directly evaluated.

As presented in Figure 2F, stromal cells from wildtype mice were sorted, and mixed with tumor cells. Cells not expressing CD31 suppressed growth, while the remaining CD31^+^ cells did not affect the tumors (Figure 2G, sorted cells shown as title for each graph). CD44 was not a discriminator as both CD44^+^ and CD44^-^ cells inhibited growth. Similarly, both the expression of Sca1 and its absence showed the same degree of growth suppression (Figure 2G). We then focused on CD31^-^ cells, and combined these with FAP and CD105. While FAP in combination with CD31^-^ failed to show enhancement of growth suppression depending on FAP expression, the combined population of CD31^-^CD105^-^ diminished growth further (Figure 2H).

In summary, a stromal subpopulation that does not express CD31 in combination with a lack of CD105 (CD31^-^CD105^-^) represents an inhibitory cells type.

### Relevance of CD105/ENDOGLIN in cancer patients

In view of the suppression of growth in the absence of CD31 and the even stronger suppression in the absence of CD105, we evaluated both CD31 and CD105 mRNA expression in two patient cohorts (GDC data portal, see methods for cohort selection). Surprisingly, the endothelial marker CD31/*PECAM1* showed no relationship to survival in melanoma or breast cancer patients, while CD105/*ENDOGLIN* mRNA expression correlated negatively with the number of days until death in melanoma patients (Figure 2I). Thus, patients that show low CD105/*ENDOGLIN* mRNA expression in their tumors (in the lowest tertile) exhibited longer survival than those in the highest tertile (Figure 2I). In breast cancer patients, the correlation was not significant (p=0.11), and only a trend that was not statistically significant was found when comparing the lowest and highest tertiles (p=0.08).

Thus, low CD105/*ENDOGLIN* expression is beneficial in melanoma patients.

### CS1, a fragment of fibronectin, enhances inhibition by stromal cells

While fibronectin pretreatment enhances growth suppression by stromal cells, deletion of fibronectin in a subpopulation of bone marrow cells counteracts the inhibition increasing cancer growth (Figure 1G and 1K). This effect, however, was only partially mediated by the classical fibronectin receptor α5β1, which binds to the amino acid sequence arginine-glycine-aspartic acid (RGD)^27^ (Figure 2E and 2F). We therefore aimed to evaluate fragments of fibronectin that attach to other receptors. CS1 represents part of the variable region normally found in plasma fibronectin (used in 1G), affects tumor development^29^, and binds to α4β1 and α4β7 integrins^30,31^.

Interestingly, pretreatment of stromal cells with CS1 suppressed growth even more than those treated with the scrambled peptide, as was seen with fibronectin (Figure 2J-K). The inhibitory effect of CS1 is not attributable to a carry-over effect of CS1 from the stromal to the tumor cells, because B16 tumor cells exposed to CS1 increased proliferation and diminished apoptosis (Supplementary-Figure 4A-B). These two effects would have resulted in the opposite effect, which is increased tumor growth. Stromal cells treated with CS1 showed no change in proliferation or apoptosis (Supplementary-Figure 4C-D).

The additional suppression by CS1 was not mediated by T cells, because repeating the experiment in the *foxn1^nu/nu^* model, in which T cells do not mature^19^ shows that growth inhibition remained (Figure 2L).

Thus, CS1 pretreatment enhances tumor growth suppression by stromal cells independent of T-cells.

### Ly6G cells mediate the suppression by stromal cells

We next sought to determine the reason for decreased growth in the presence of stromal cells. Because a role for T cells in mediating growth suppression was excluded (Figure 1D and 2L), we focused on myeloid cells.

The percentage of immune cells was evaluated in the presence of stromal cells without and with CS1 pretreatment (Supplementary-Figure 5A). Ly6G^+^ granulocytic cells changed in the opposite direction to tumor size and both correlated negatively, as if Ly6G^+^ cells mediated the suppression of growth (Figure 3A). Similarly, a higher percentage of Ly6G^+^ cells was observed in the presence of stromal cells (Figure 3B), when growth was suppressed with pretreatment with plasma fibronectin (pFN), and in the transgenic mouse models with diminished growth (white bars) (Figure 3C). Notably, the combination of CD11b^+^Ly6G^+^Ly6C^+^, which may reflect a population of myeloid-derived suppressor cells (MDSCs), also increased in the presence of stromal cells, that is, in small tumors (Figure 3D)^19^.

**Figure 3.**
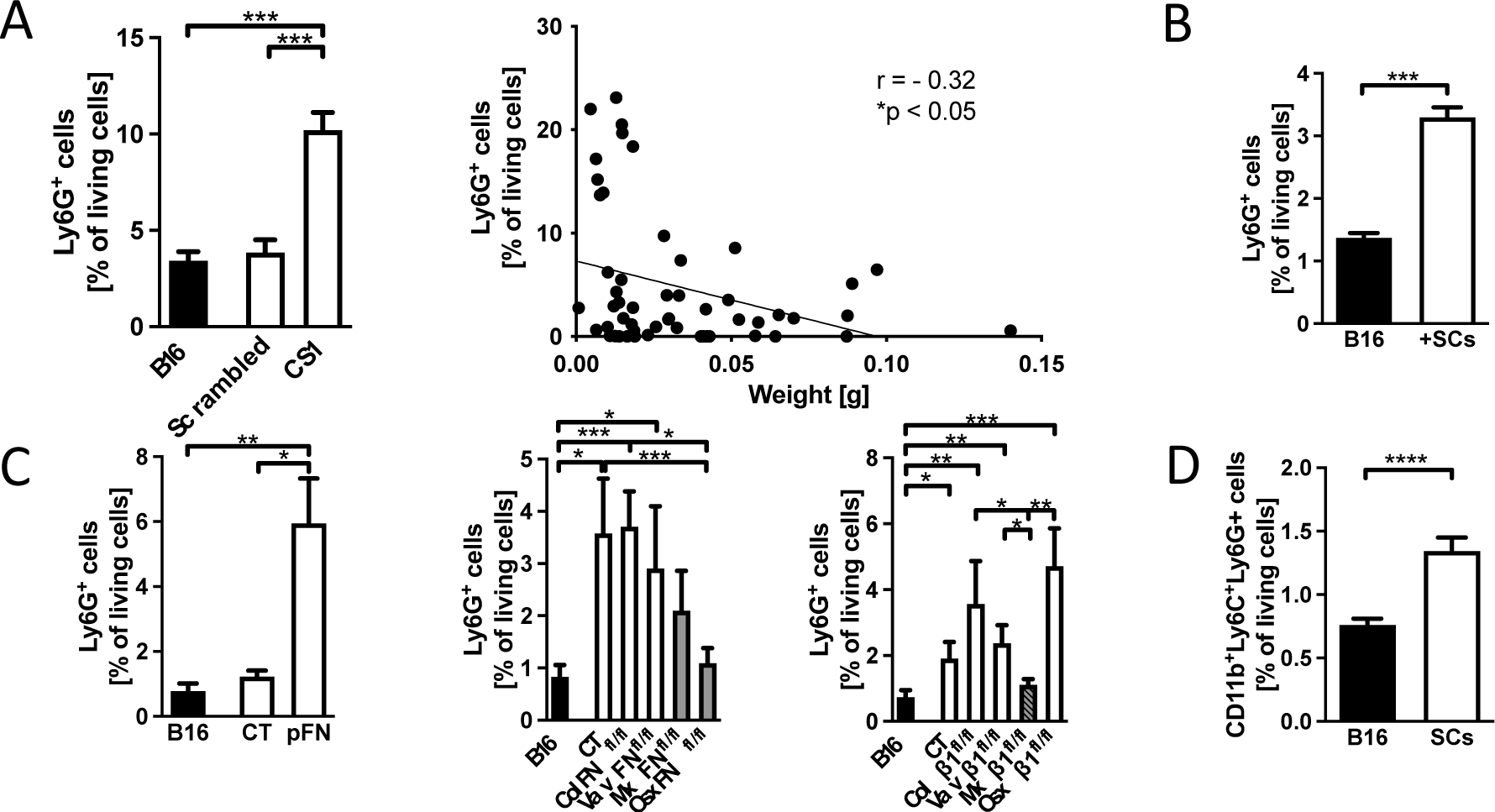
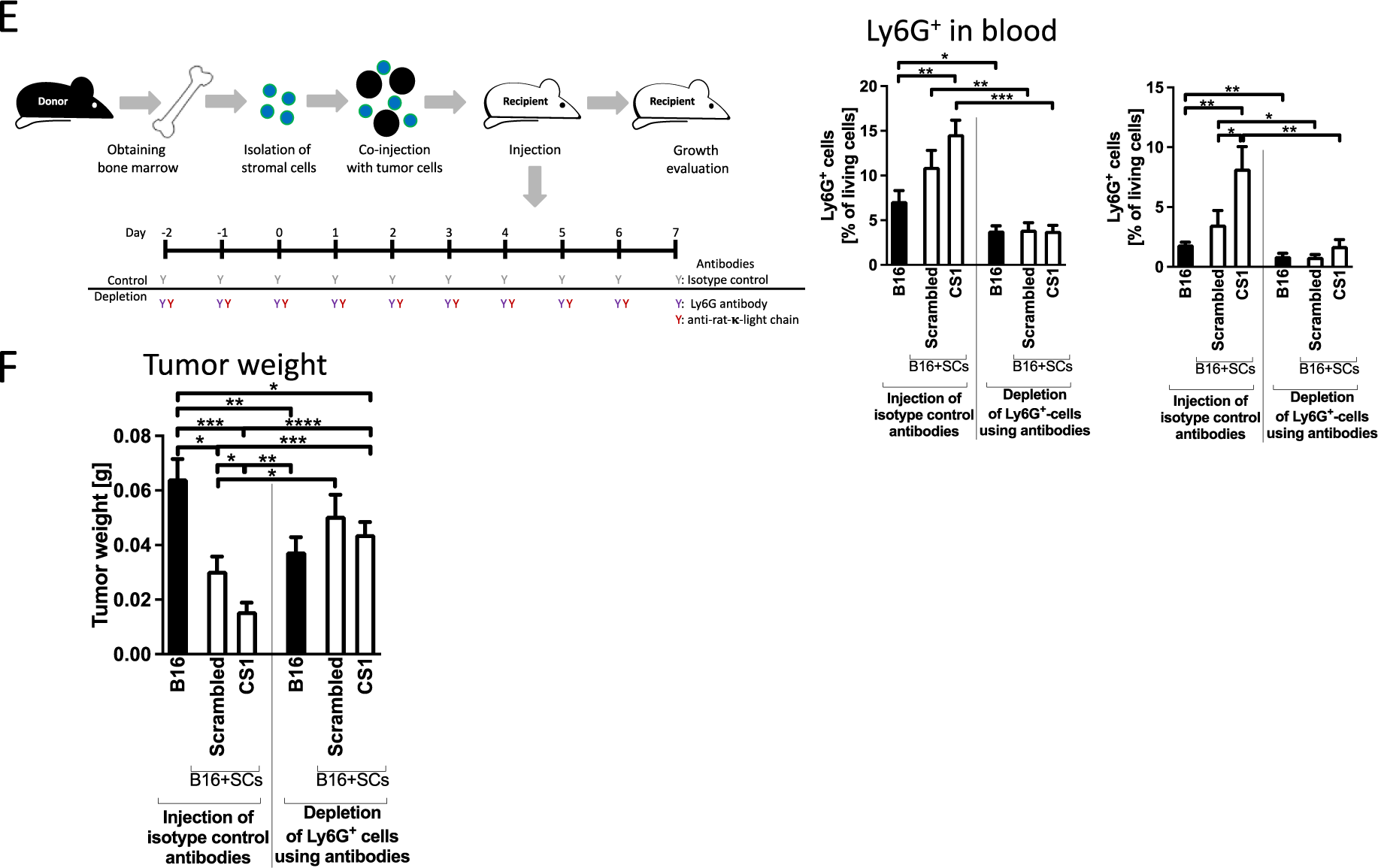
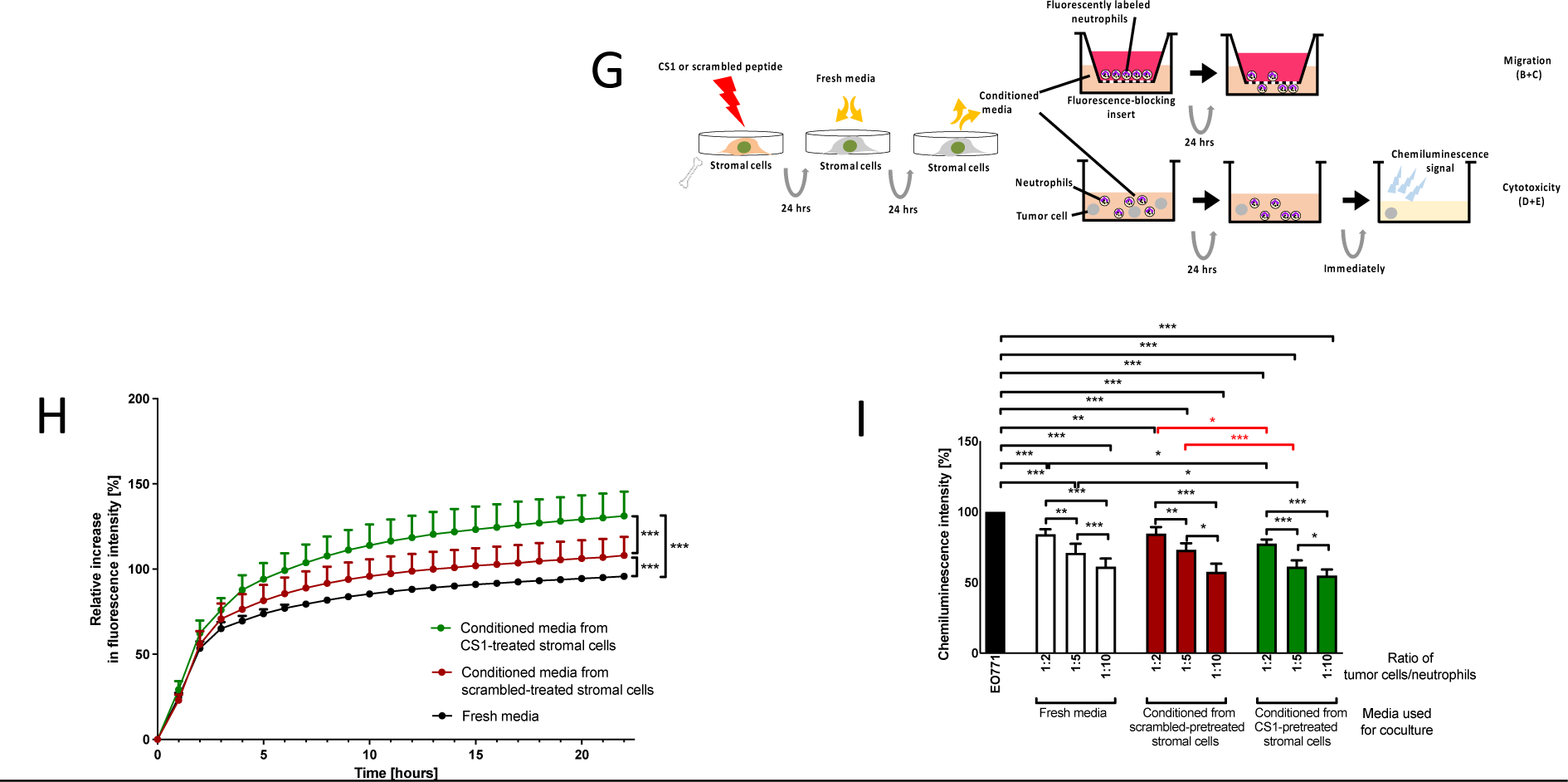
A-D. Ly6G^+^ cells are increased in the presence of inhibitory stromal cells. Ly6G^+^ cells mediate growth suppression by stromal cells G-I. CS1 enhances Ly6G+ cell migration and affects activity in vitro. A. The percentage of Ly6G-expressing cells in increased in tumors containing CS1-pretreated stromal cell. N=12/10/11, *p<0.05, ***p<0.005. Correlation between tumor weight and Ly6G^+^ cells is negative, with higher percentages of Ly6G^+^ cells in smaller tumors. Pearson’s r correlation is shown. *p<0.05, N=55 and includes tumors from mice injected with B16 without stromal cells, or pretreated stromal cells exposed to scrambled peptide or CS1. B-C. Changes in Ly6G^+^ cells in the presence of freshly isolated stromal cells (B), pretreatment of stromal cells with plasma fibronectin, transgenic mouse models with fibronectin deletion or β1 integrin deletion (C). Columns in grey show the genotype without or with incomplete growth suppression. Two exceptions to the higher percentages of Ly6G^+^ in smaller tumors are limited to the control groups for CS1 and pFN (left graphs in Figure 3A and 3C), in which cultured stromal cells were injected as opposed to Figure 3B using fresh stromal cells. The tumor experiments were presented in Figure 1B (presence of stromal cells), 1G (pretreatment with plasma fibronectin: pFN), 1K (fibronectin transgenic models) and 2E (β1 integrin transgenic models). Freshly isolated stromal cells: N=12/12, pFN pretreatment: N=6/7/9; fibronectin deletion: N=13/14/6/8/6/8, β1 integrin deletion: N=17/16/9/12/15/11, *p<0.05, **p<0.01, ***p<0.001. D. Myeloid-derived suppressor cells (MDSCs: CD11b^+^Ly6G^+^Ly6C^+^) are increased in the presence of stromal cells. N=12/12. ****p<0.0001. Comparisons were performed using t-tests or ANOVA followed by t-tests. E. Schematic showing the design of the experiments: isolated stromal cells (SCs) were mixed with melanoma B16 cells and injected into mice. 2 days before injecting the cells and then daily until day 6 after cell injection, either isotype controls or an antibody directed against Ly6G were administered subcutaneously. In addition, on alternate days starting two days before tumor cell injection an anti-rat-κ-light chain antibody was administered to improve efficacy of depletion. On day 7, animals were euthanized. Graphs show successful depletion of Ly6G^+^ cells in peripheral blood and in tumors as confirmed by flow cytometry at the time of euthanasia. ANOVA was significant and followed by pair comparisons using t-tests. N=10/12/12/10/12/12. *p<0.05, **p<0.01, ***p<0.005. F. Depletion of Ly6G^+^ cells resulted in loss of the suppressive effect of stromal cells as well as loss of the additional inhibitory effect of CS1 (Two white bars on the right part of the graph). B16 tumors without stromal cells were smaller after depletion (black bars). If ANOVA was significant, pair comparisons using t-tests were performed. N=10/12/12/10/12/12. *p<0.05, **p<0.01. G. Schematic showing experimental design to evaluate migration and cytotoxicity of Ly6G^+^ cells. Freshly isolated stromal cells were pretreated with CS1. 24 hours later, fresh medium was added for another 24 hours. Conditioned media were applied in the lower well in a transwell assay in the migration assay and migration of fluorescently labeled neutrophils through the floor of the insert was evaluated by measuring the fluorescence in the lower well. Conditioned media were also added to the mixture of luciferase-expressing tumor cells and freshly isolated Ly6G^+^ cells at different ratios. 24 hours later, the luminescence signal of the remaining tumor cells was evaluated by adding luciferin. H. CS1 enhances migration of Ly6G^+^ cells towards conditioned media from CS1-pretreated stromal cells (in green) compared to scrambled-pretreated cells (in red). N=11/11/11; ***p<0.001. Non-linear regression was performed. I. Cytotoxicity assay using Ly6G^+^ cells isolated from the bone marrow and cocultured with EO771/luc^+^ tumor cells in the presence of conditioned media shows statistical differences. The comparisons in red show that the amount of cancer cells differed between CS1-(green bars) and scrambled-pretreated conditioned media (red bars). This suggests higher killing efficacy of Ly6G^+^ cells, but this effect is small. Note that less remaining tumor cells results in lower luminescence signal intensity and is compatible with smaller tumors. N= 18/18/18/18/18/18/18/18/18/18, *p<0.05, **p<0.01, ***p<0.001. ANOVA, if significant, was followed by t-test comparisons.

To establish a causal relationship between the changes in Ly6G and tumor suppression, we depleted Ly6G^+^ cells using an established protocol (Figure 3E)^32^, and confirmed depletion in in the peripheral blood and in the tumors (Figure 3E). In the absence of Ly6G^+^ cells, stromal cells no longer resulted in growth inhibition (two white bars on the right in Figure 3F). Interestingly, there was a concomitant decrease in growth of B16 in the absence of stromal cells and Ly6G^+^ cells (black bars in Figure 3F).

This shows that in the absence of stromal cells, cells that express Ly6G^+^ support tumor growth; however, in the presence of stromal cells, inhibition of growth is mediated by Ly6G^+^ cells.

### CS1-treated stromal cells support migration of Ly6G^+^ cells

To determine whether Ly6G^+^ cells diminished growth due to their increased number or enhanced tumor-cell killing effects in the presence of stromal cells, *in vitro* experiments were performed.

The migration of Ly6G^+^ cells was evaluated. Stromal cells were pretreated with CS1 for 24 hours, and fresh medium was added for 24 hours. The conditioned media were then added to the bottom wells of a transwell assay (Figure 3G). Ly6G^+^ cells migration towards the conditioned medium was enhanced when the media originated from CS1-pretreated cells (Figure 3H). This was not the case when using the fibroblastic cell line NIH3T3 (Supplementary-Figure 5C). Thus, CS1 pretreatment may enhance the recruitment of Ly6G^+^ cells to the tumor.

In a cytotoxicity assay, Ly6G^+^ cells isolated from the bone marrow were cocultured with tumor cells, which express luciferase (/luc^+^). Co-culturing the cells in conditioned media from CS1-pretreated stromal cells resulted in less luminescence signal, in line with the loss of tumor cells (Figure 3I and Supplementary-Figure 5E, red comparisons). Even though the effect is small, it is possible that a biologically relevant effect of the stromal cells in enhancing cytotoxic Ly6G^+^ cell activity *in vivo* takes place.

### CS1 effects are mediated by TLR4 in vitro

Immune cell migration is usually mediated by cytokines. Stromal cells treated with pFN or CS1 showed increased mRNA expression of several cytokines (CXCL1, CXCL2, TNF-α, IL-1β and IFNβ), which enhance Ly6G^+^ recruitment, activity, and/or polarization (Figure 4A)^11,33^. These cytokines are usually produced in response to the activation of NF-κB signaling^33^. We therefore evaluated and confirmed NF-κB translocation into the nucleus in response to plasma fibronectin (pFN) and CS1 treatment (Figure 4B). IκBα and NF-κB phosphorylation increased in line with the activation of this signaling cascade (Supplementary-Figure 6A-B). Because pFN and CS1 can bind to integrins^30^, molecules downstream of integrin signaling were examined. Both pFN and CS1 induced the phosphorylation of focal adhesion kinase, AKT, and ERK (Figure 4C) ^30,34^. These data confirmed activation of NF-κB and integrin signaling.

**Figure 4.**
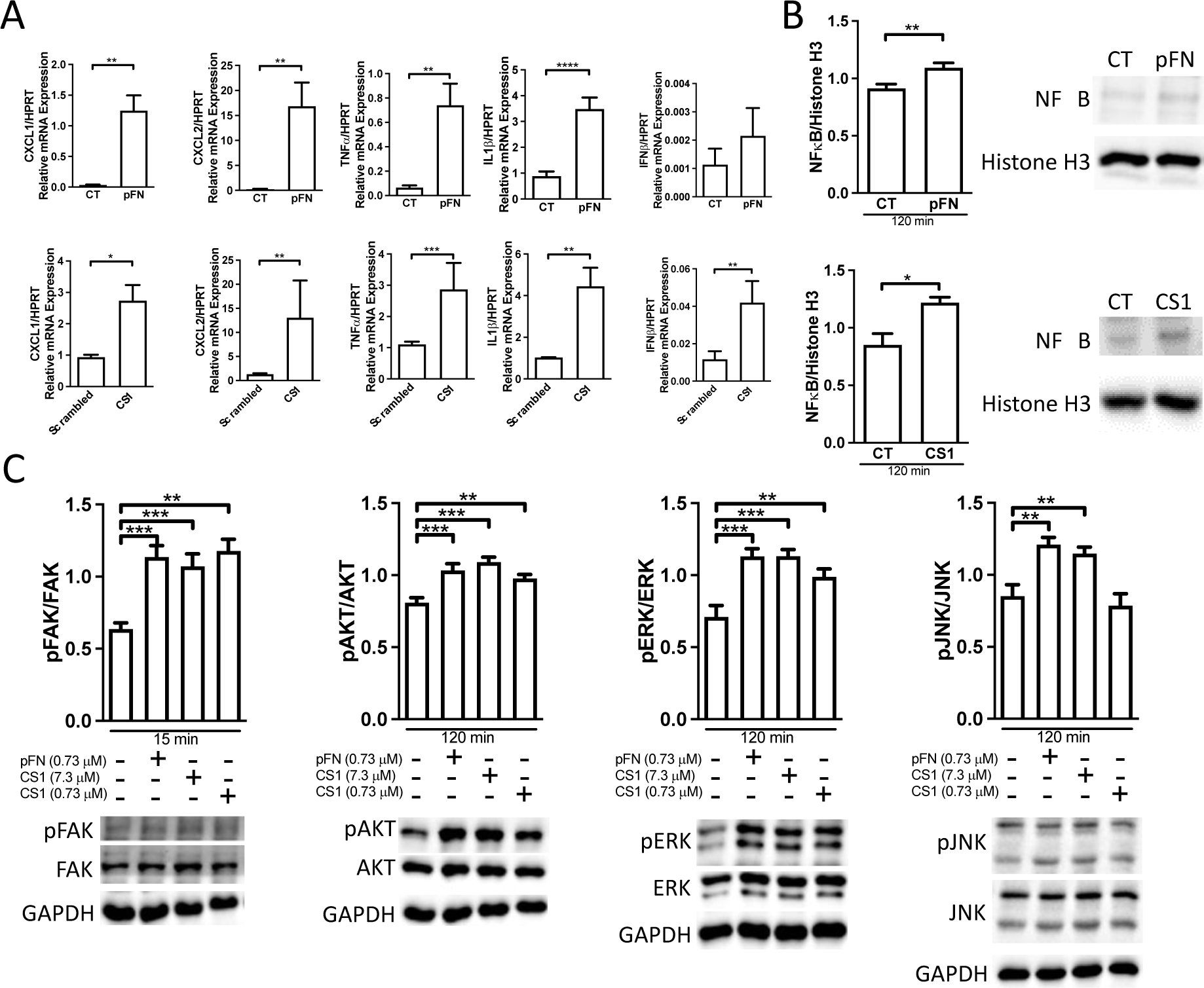
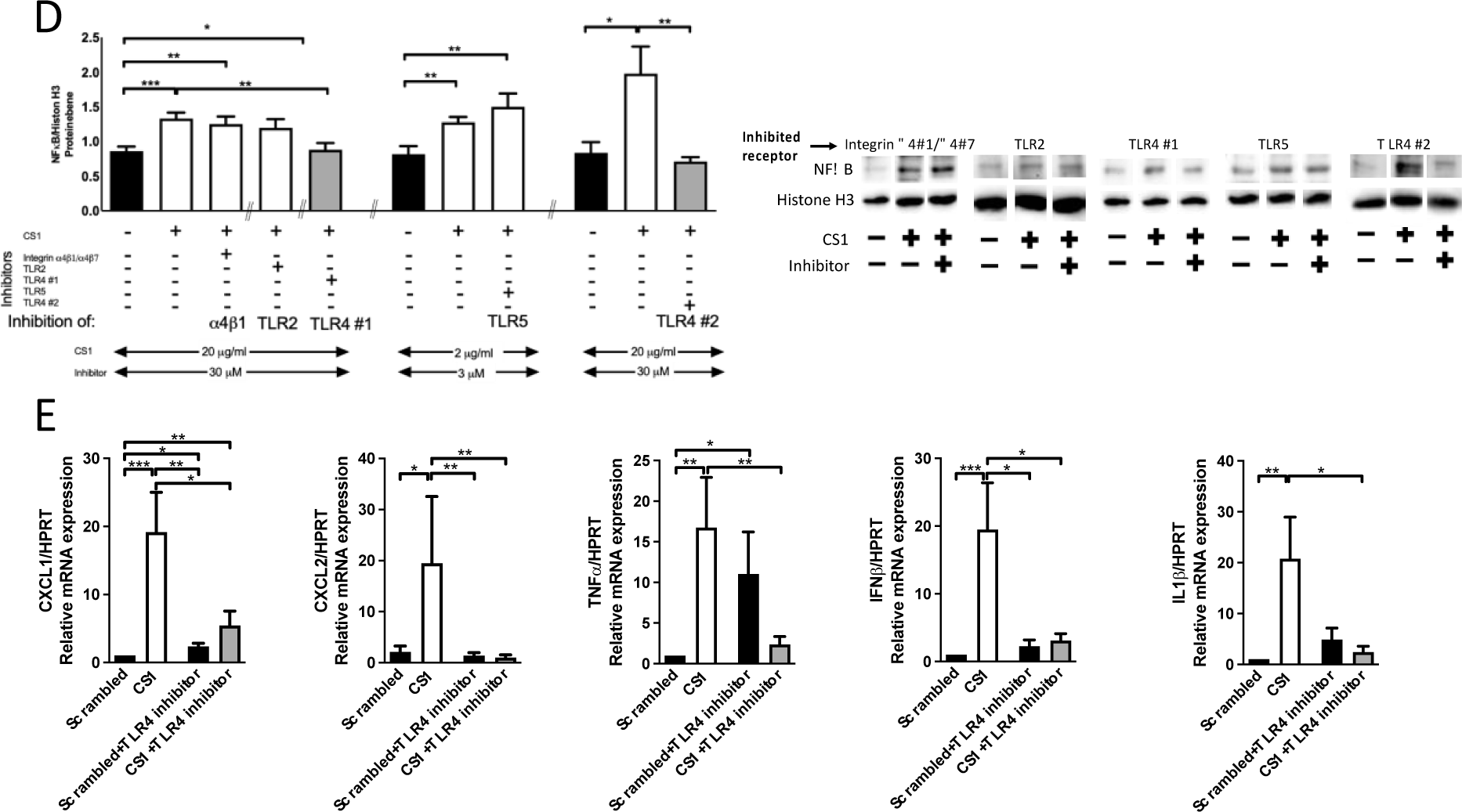
Evaluating cytokines and signaling TLR4 mediates CS1 effects on stromal cells in vitro. **A.** Increased cytokine mRNA expression by stromal cells treated with plasma fibronectin (pFN) or CS1. mRNA expression of cytokines known to enhance Ly6G migration, activation or polarization was evaluated 24 hours after treatment of freshly isolated stromal cells with plasma fibronectin (pFN) 160 μg/ml (0.73 μM) or CS1 20 μg/ml (7.3 μM). Pairs were evaluated using non-parametric t-tests. For pFN: CXCL1: N=9/18, CXCL2: N=11/13, TNFα: N=14/17, IFNβ: N=6/8, IL1β: N=14/15. For CS1: CXCL1: N=17/14, CXCL2: N=9/10, TNFα: N=13/15, IFNβ: N=22/17, IL1β: N=14/11. *p<0.05, **p<0.01, ***p<0.001. **B. Activation of NF-κB-mediated signal pathway.** NF-κB translocation into the nucleus of stromal cells was confirmed by adding plasma fibronectin or CS1 to stromal cells for 120 minutes, generating nuclear extracts and examining the amount of NF-κB in the nucleus. N=10/10 for fibronectin and N=4/4 for CS1, *p<0.05. Data evaluated by t-test. **C. Changes in integrin signaling.** Focal adhesion kinase phosphorylation (pFAK) increased 15 minutes after pFN or CS1 addition to stromal cells. N= 4/4/4/4, ***p<0.001. Phosphorylation of AKT, ERK and JNK is shown for the time point of 2 hours after addition of pFN or CS1. pAKT increased after exposure to the additives. N=10/10/10/10, **p<0.01, ***p<0.001. Phosphorylation of ERK increased too. N=10/10/10/10, **p<0.01, ***p<0.001. Phosphorylation of JNK in response to additives is not increased with the low concentration of CS1. N=9/9/9/9, **p<0.01. ANOVA was performed and, if significant, followed by t-tests. Freshly isolated stromal cells were starved overnight, exposed to 160 µg/ml plasma fibronectin and 20 µg/ml and 2 µg/ml of CS1 for different times and cell lysates were collected and evaluated by western blotting. The time points presented are written below the x-axis. D. Only TLR4 inhibition prevented NF-κB translocation shown in grey compared to control (CT) shown in black. The inhibition was confirmed using a second TLR4 inhibitor shown on the far right in grey compared to CT in black. ANOVA was performed and followed by t-tests. N=23/23/21/11/14/8/8/8/9/9/10, *p<0.05, **p<0.01, ***p<0.001. Stromal cells were exposed to the inhibitors BIO5192 (for integrin α4β1 and α4β7), TLR2-IN-C29 (TLR2), TLR4-IN-C34 (TLR4#1) and TAK-242 (TLR4#2) at a concentration of 30 µM and TH1020 (TLR5) at a concentration of 3 µM for 1 hour prior to treatment with 20 μg/ml CS1 (2 µg/ml in conjunction with TH1020) for an additional hour. The nuclear lysates were evaluated by western blotting for NF-κB translocation. Examples from translocation experiments are shown for the use of the inhibitors in the presence of CS1. E. TLR4 inhibition normalized the cytokines that were increased in response to CS1 treatment. CXCL1: N=15/11/11/13, CXCL2: N=16/11/14/14, TNFα: N=11/11/10/10, IFNβ: N=14/11/11/13, IL1β: N=11/12/9/10, IL6: N=13/12/9/13. *p<0.05, **p<0.01, ***p<0.001. Stromal cells were exposed to the inhibitors (at the concentrations outlined in D) for 1 hour prior to treatment with 20 μg/ml CS1 for 24 hours. mRNA expression of the cytokines was determined by qPCR. ANOVA was performed and followed by t-tests in D and E.

We next aimed to determine which receptor mediates NF-κB translocation to the nucleus in response to pFN and CS1. Various domains of fibronectin activate one or more of the following receptors: Toll-like receptor 2 (TLR2), TLR4, and TLR5 as well as integrin α4β1 and α4β7^35,36^, which can stimulate NF-κB translocation or interact with NF-κB signaling^33^. These receptors, except β7, were expressed on the cell surface of stromal cells (Supplementary-Figure 6C). NF-kB translocation was only prevented after TLR4 inhibition (using two different chemical inhibitors) (Figure 4D). Importantly, pretreatment with a TLR4 inhibitor for one hour before CS1 stimulation prevented changes in cytokine mRNA expression (Figure 4E).

Thus, TLR4 on stromal cells is a potential mediator of CS1 effects.

### TLR4-signaling mediates the immune response of stromal cells against cancer

In order to define the relevance of TLR4 expression on stromal cells, we sorted TLR4^+^ and TLR4^-^ stromal cells (Figure 2F). No difference was seen between the effects of the two cell populations (Figure 5A). We then inhibited TLR4 in order to determine whether CS1 effects *in vivo* are mediated by TLR4 activation in stromal cells. Stromal cells were first exposed to a TLR4 inhibitor for one hour and then to CS1 before injection in mice (Figure 5B). As suggested by the *in vitro* experiments in Figure 4D, TLR4 inhibition prevented the increase in Ly6G^+^ cells in response to CS1 (compare the two right columns with the two middle columns in Figure 5C). Suppression of growth in response to CS1 was no longer detectable (again, compare the two right columns with the two middle columns in Figure 5D). Interestingly, TLR4 inhibition failed to completely restore the suppressed growth in the presence of stromal cells.

**Figure 5.**
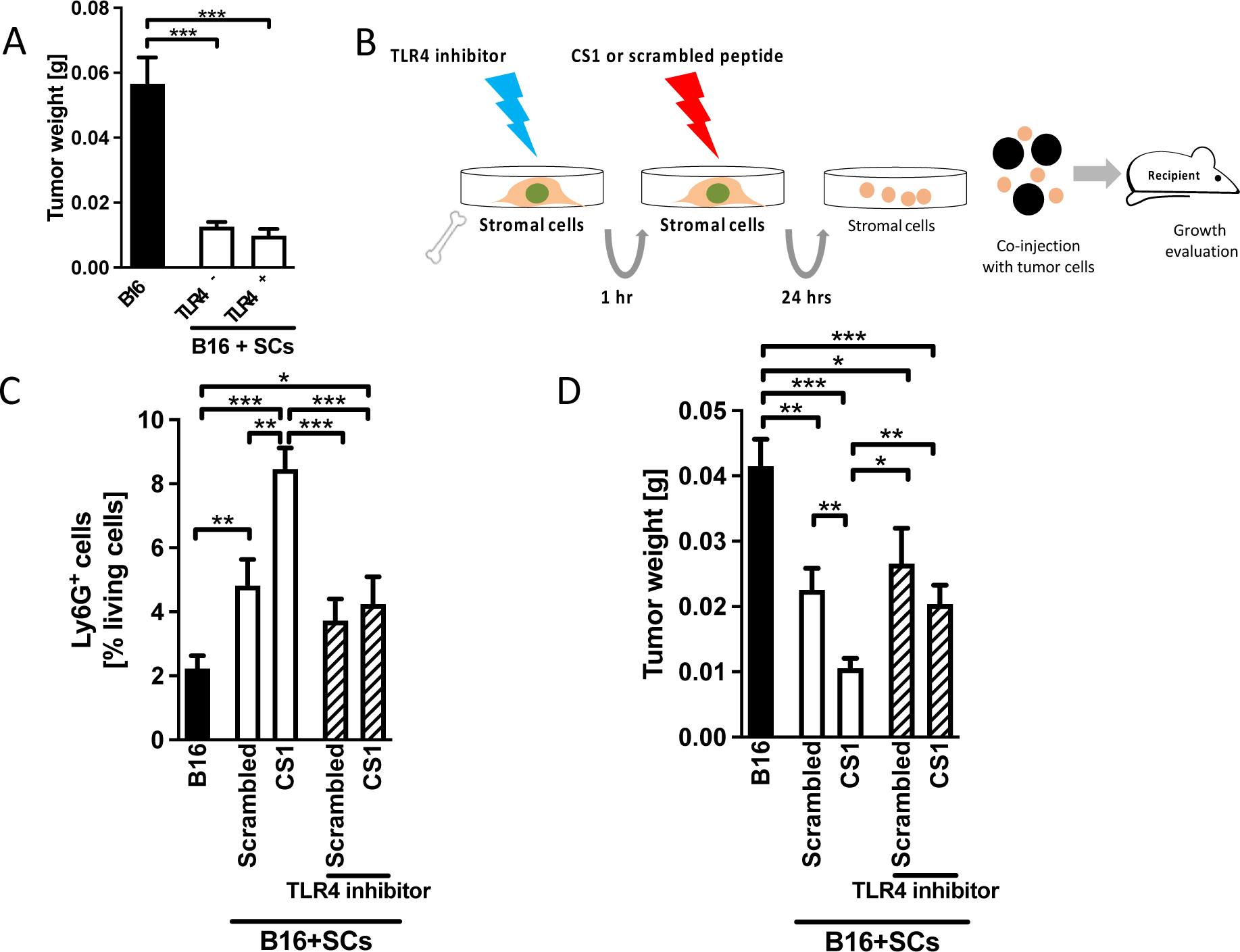
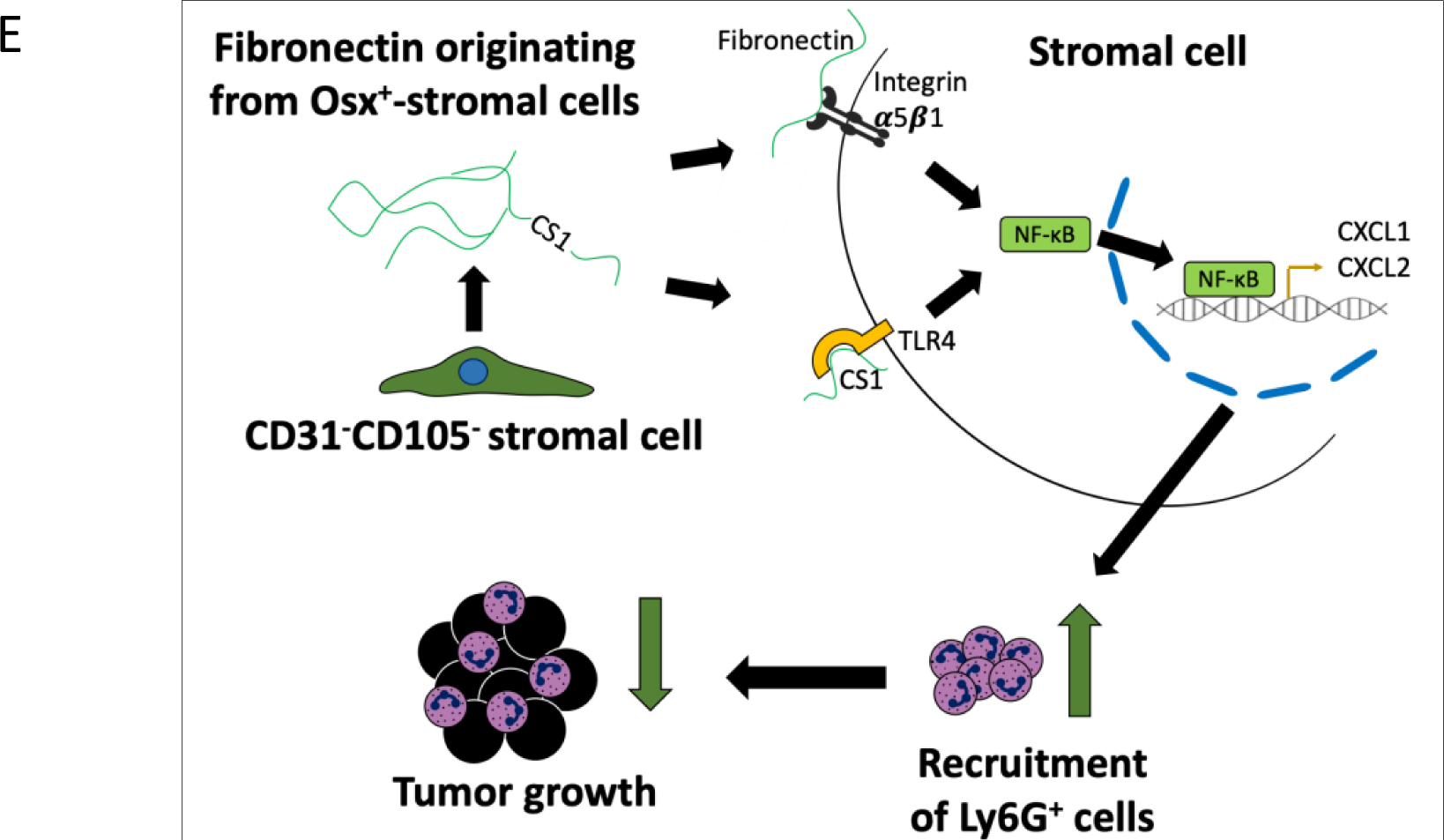
A-D. TLR4 mediates CS1 effects on stromal cells in vivo E. Summary Bone marrow stromal cells inhibit cancer growth through fibronectin actions. A. No difference on growth inhibition between stromal cells (SCs) that express TLR4 (TLR4^+^) and those that do not (TLR4^-^). The experiment was performed as shown in 2F and 3G. Bone marrow stromal cells were sorted based on TLR4 expression and injected. Differences were examined by t-tests. N=6/11/6. **p<0.01, ***p<0.001, ****p<0.0001. B. Schematic showing the pretreatment of isolated stromal cells with TLR4 inhibitor TLR4-IN-C34 at 30 μM for 1 hour, media were changed and CS1 added at 20 μg/ml for 24 hours. Stromal cells were then collected, mixed with B16 melanoma cells and injected subcutaneously. C. TLR4 Inhibition prevents CS1-mediated increase in the percentage of Ly6G^+^ cells (last bar). N=12/12/12/12/12. *p<0.05, **p<0.01, ***p<0.001. D. TLR4 Inhibition prevents suppression of tumor size by stromal cells pretreated with CS1. Tumor size is restored to the level of scrambled-peptide-pretreated stromal cells and is larger than the size in response to CS1 pretreatment only. Cells were treated as shown in B. N=12/12/12/12/12. *p<0.05, **p<0.01, ***p<0.001. Analysis by ANOVA was followed by t-tests if significant. Stromal cells that strongly inhibit tumor growth are characterized by the combined absence of CD31/pecam1 and CD105/endoglin. Mechanistically, fibronectin produced by a small subpopulation expressing osterix/sp7 acts directly on its receptor, α5β1 integrin, or via CS1 on TLR4 located on stromal cells. This enhances NF-κB translocation, leading to the production of chemokines such as CXCL1. Consequently, there is increased influx of Ly6G^+^ cells, leading to immune-mediated inhibition of early tumor growth.

In summary, TLR4 mediates increased neutrophils and inhibition of tumor growth in the presence of CS1.

## Discussion

We show that bone marrow stromal cells that do not express CD31 and CD105 are able to suppress tumor growth, and the subpopulation that expresses osterix/sp7 produces fibronectin that mediates this inhibition *in vivo*. Indeed, activating bone marrow stromal cells with fibronectin or the fragment CS1 enhances NF-κB signaling and the production of cytokines such as CXCL1 and CXCL2, which then boosts recruitment of immune cells that express Ly6G. These immune cells suppress tumor growth (Figure 5E).

It is surprising that only a few co-injected stromal cells remained in the tumor after three days (Figure 1F), and despite this, growth suppression was found after two weeks. A possible explanation is that the presence of the fibroblastic cells at the time of injection induces a delay in growth that is large enough to affect the outcome at 2 weeks. In support of this, in a previous experiment, that lasted for 6 weeks, the tumor remained smaller^16^. The initial stimulus thus seems to increase Ly6G^+^ recruitment/activation and could directly or through the Ly6G^+^ cells attract more cells and suppress tumor growth^37^.

Fibronectin is not only produced *in vivo* in mammals by a large variety of cells but its supportive role in cancer has been documented ^18,20^. It was therefore surprising, that both fibronectin and CS1 suppressed growth further and suggests a new role for fibronectin, depending on its source. A related pattern was shown for tenascin C. On one hand, tumoral tenascin C drives tumor progression by polarizing macrophages into the M2 phenotype, while, on the other hand, host-derived tenascin C (presumably from stromal and/or immune cells) polarizes the macrophages into the M1 phenotype^38^.

The inhibitory effect of fibronectin is at least partially mediated by fibronectin-binding α5β1 integrin and that of CS1 by TLR4. Indeed, a TLR4 agonist was found to diminish cancer growth^39^. TLR4 can have opposing effects, however, depending on the context and may confer a survival advantage for the tumor cells^40^. Since TLR4 is normally expressed on immune cells^41^, it seems that with regard to this function, fibroblasts took over some immune cell characteristics. This is supported by the production of cytokines and the findings on signaling shown in Figure 4.

The inhibitory stromal population contains a subgroup (4% of stromal cells) that activates sp7/osterix and hence is related to osteoblast progenitors. It differs from the remaining stromal cells in its proteomic signature (Supplementary-Table 1) suggesting that it represents a distinct group that would be beneficial if present in the tumor. Clinical data analysis failed to detect a change in sp7 mRNA expression. Instead, in line with the characterization of CD31^-^CD105^-^ as inhibitory cells, low CD105/*ENDOGLIN* mRNA expression in melanoma patients was associated with increased survival (Figure 2I). Our results are in line with work showing that in pancreas cancer, CD105^-^ fibroblasts suppress tumor progression^10^, and high expression is associated with more metastases^42^. Furthermore, since both molecules are expressed on endothelial cells and support angiogenesis, it is tempting to speculate that stromal cells exert an inhibitory effect unless they can support angiogenesis.

It seems contradictive that fibronectin supports cancer growth unless it originates from the stromal cells. In view of the findings in patients, one possible explanation is that it needs to originate from stromal cells that do not express CD105, which then directly or indirectly recruit Ly6G^+^ cells. Neutrophils modulate tumor cells through a variety of mechanisms^11,43^. In the presence of stromal cells, Ly6G^+^ cells suppressed growth making them N1-like (Figure 3F)^44^. The opposite to a report on macrophages, where fibroblasts can change the characteristics of macrophages to become M2 pro-tumorigenic^9^. Although there is evidence that the number of neutrophils increases only in the first 7 days after injection^45^, our data showed that the increase can last up to two weeks (For example Figure 3B). Suppression of growth could be mostly mediated by the larger numbers. It is nevertheless possible that the Ly6G^+^ cells, once exposed to stromal cells, enhance their anti-tumor activity *in vivo*. In support of this are the findings, albeit small, on cytotoxicity *in vitro* (Figure 3I). A dual action (migration and activation) of neutrophils was also reported in other settings^12^.

In summary, we characterized inhibitory stromal subpopulations and established that fibronectin acts on stromal cell receptors to increase the production of chemotactic cytokines and enhance recruitment of anti-tumorigenic neutrophils leading to tumor suppression. This does not require functional T cells. Evaluating how to take advantage of this mechanism in early cancer might be a worthwhile endeavor.

## Material and methods

### 1. Patient data

The cohorts for melanoma (193 patients) and breast cancer (135 patients) were generated from the Genomic data commons (GDC) data portal by restricting the cohort to patients in whom the primary tumor was located in the skin (for melanoma) or in the breast, with no prior malignancy, prior treatment, or synchronous malignancy. Only patients who had died and in whom the days to death were reported were included. Preliminary analysis was performed directly on the website under gene expression clustering followed by downloading the original data for statistical analysis.

### 2. Mice and experimental procedures

C57BL/6 mice were obtained from Janvier Laboratories (RRID:MGI:2159769). Immune competent and T cell deficient CD1-Foxn1^nu^ mice were also used (RRID:MGI:5522729)^46^. The former were used to evaluate the impact of the genetic background of experimental animals as well as for the conditional fibronectin and integrin β1 knockout experiments while the latter allowed us to elucidate the lack of T cells in our models. The mice injected with β16 cells were both sexes and with EO5771 were females. All were aged 4-6 weeks. Each experiment included a control group receiving only tumor cells. Since the global knockout for fibronectin or β1 integrin lead to embryonic lethality, animals homozygous for the floxed fibronectin or β1 integrin gene were used (RRID:IMSR_JAX:029624 and RRID:MGI:4358370)^22,25^. The conditional knockout animals had a floxed fibronectin or β1 integrin gene and carried the Colα(1)I, Mx1, Vav, or osterix/sp7 (Osx) promotor driving cre-recombinase^22,23,25,47^. Littermate controls not carrying the cre-recombinase transgene were included as appropriate. To evaluate the activity of the promoters used, mice expressing tdTomato (JAX #007909) were mated with mice carrying the respective promotors without any floxed genes^22^. For some experiments, tdTomato was introduced into β1^fl/fl^ mice. Animal studies followed international including EU, national, and institutional guidelines for humane animal treatment, complied with the ARRIVE guidelines and relevant legislation, and were approved by the appropriate office for animal welfare in the state of Baden-Wuerttemberg, Germany (Regierungspraesidium Karlsruhe). The protocols used carry the following numbers: T-44/20, G-21/18, G-249/18, G-242/19, G-5/21, G-102/21, G-216/21, G-219/21, G-222/21, G-230/21, G-277/21, G-280/21, G-283/21 and G-284/21. The sample size for each experiment was determined based on power calculations relying on experimental evidence from previous experiments or *in vitro* studies. The number of mice in each group is mentioned in the figure legend. Animals in which tumor did not grow were excluded. The mice were randomly assigned to the various groups. At the time of killing, animals were selected randomly and parameters evaluated without knowledge of the group assignment. Outcome measures included tumor weight, tumor volume and flow cytometry results (see below).

B16 melanomas were induced by injection of 10^6^ tumor cells together with either 2×10^6^, 1×10^6^ or 0.1×10^6^ bone marrow stromal cells subcutaneously in the right flank of the animals. Mice were sacrificed 7 or 14 days after tumor cell injection.

Orthotopic EO771 mammary tumors was induced by injection of 10^6^ tumor cells together with 10^5^ bone marrow stromal cells into the left abdominal mammary gland. Mice were sacrificed after 14 days.

Depletion of Ly6G-expressing cells started two days prior to tumor induction and was performed by daily intraperitoneal injection of 50 µg anti-mouse Ly6G antibodies (#127650, Biolegend, RRID:AB_2572001) in conjunction with 50 µg of an anti-rat κ-light chain antibody every other day (#BE0122, Bio X Cell, RRID:AB_10951292) to enhance depletion efficiency. Control mice received 50 µg of an IgG2a κ isotype antibody daily (#BE0089, Bio X Cell, RRID:AB_1107769). The mice were sacrificed seven days after tumor injection. To confirm Ly6G depletion blood was collected and red blood cells were lysed. The obtained immune cells were then fixed in 1% PFA and permeabilized with 0.1% triton X. This was necessary to stain extra- and intracellular Ly6G to counteract the possibility of overlooking cells masked by the anti-mouse Ly6G injection. The permeabilized cells were then blocked with 5% BSA for 15 minutes and stained for flow cytometry.

The tumors were isolated and digested for one hour in FCS-free DMEM (PAN Biotech) containing 1 µg/ml collagenase (Nordmark Biochemicals) and 0.5 µl/ml DNase (Qiagen).

### 3. Cell isolation and culture

#### Stromal cell isolation

Flushed bone marrow was depleted using magnetic protein-G Dynabeads (Invitrogen), which bind mammalian antibodies^22^. Briefly, the magnetic beads (50μl/10^7^ cells) were incubated with 25 μl unconjugated anti-mouse CD45 antibody ((#103102, Biolegend, RRID:AB_312967) in 250 μl D-PBS for 30 minutes while shaking. Through use of a magnet, excess antibody was removed by washing twice with PBS. The bone marrow cells (10^7^) were then exposed to the precoated beads for 30 minutes while shaking to bind the CD45^+^ cells to the beads. The supernatant was collected and contained CD45^-^ stromal cells.

CD45^+^ cells were isolated similarly by using sheep anti-rat Dynabeads, removing the supernatant containing the stromal cells, and separating the cells from the beads using the isolation buffer based on the manufacturer’s instructions.

#### *In vitro* treatments

Pretreatment of stromal cells was performed in 2 ml αMEM media (Gibco) per 10^7^ cells lacking FCS in 15 ml conical tubes. The peptides were added after one hour of equilibration. The cells were pretreated with plasma fibronectin (160 µg/ml), the CS1 peptide (20 µg/ml) (DELPQLVTLPHPNLHGPEILDVPST), or a scrambled peptide (20 µg/ml) (GDPELNITLSVPLPTHLQEPDPVLH) for 24 hours. The peptides were produced by the core facility at the Max-Planck Institute for Biochemistry (Martinsried, Germany). After pretreatment and before injection, the cells were washed with PBS three times.

For the pharmacological inhibition of receptors, the stromal cells were pretreated one hour before the addition of the respective peptide. The inhibitors used were BIO5192 (Tocris), TLR2-IN-C29 (Selleck Chemicals), TLR4-IN-C34 (Sigma-Aldrich), TAK-242 (Sigma-Aldrich) and TH 1020 (Tocris). All inhibitors were used at a final concentration of 30 µM except TH 1020 which was added at 3 µM (and accordingly, a ten-fold lower concentration of the CS1 peptide was used in conjunction with this inhibitor).

#### Culture of tumor cells

B16-F10, EO771, and EO771/luc^+^ cancer cells (ATCC, RRID:CVCL_F936 and CVCL_GR23) were cultured in DMEM medium (Gibco) supplemented with 10% fetal calf serum (FCS) (PAN Biotech) and 1% penicillin/streptomycin (Gibco). Mycoplasma contamination was excluded every 4-6 weeks.

#### Isolation of neutrophils

Murine neutrophils were isolated using a Histopaque density gradient (Sigma-Aldrich). Briefly, 3 ml of histopaque-1119 was pipetted into a 15 ml conical tube and 3 ml of histopaque-1077 was carefully layered on top of it. Next, 1 ml of murine bone marrow, which was isolated as described above, or murine blood was layered on top of histopaque-1077 and then centrifuged for 30 minutes at 872 g. Neutrophils were collected at the interface between histopaque-1077 and histopaque-1119. To produce conditioned media for the transwell and cytotoxicity assays, stromal cells were isolated and pretreated with the CS1 or scrambled peptide, as described above. Afterwards, αMEM supplemented with 10% FCS and 1% penicillin/streptomycin was added to generate conditioned media of the differently stimulated stromal cells. Conditioned media were collected and always used immediately.

#### Transwell assay

For the transwell assay, isolated neutrophils were fluorescently labeled with CFSE (1:1000) for 20 minutes at 37°C. In the lower compartment of the Corning® FluoroBlok™ Transwell system, 450 µL of conditioned media was mixed with 300 µL αMEM containing 10% FCS and 1% penicillin/streptomycin. The insert was then placed into the wells and loaded with FCS-free αMEM containing 10,000-15,000 CFSE-labeled neutrophils from murine blood. The increase in fluorescence intensity due to migrating cells was then measured once per hour and calculated in relation to the fluorescence intensity at 0 hours (Spark, Tecan).

#### Cytotoxicity assay

For the cytotoxicity assay, luciferase-transfected EO771/luc^+^ or MDA-MB-231-B/luc^+^ cells were plated in black 96-well plates. Neutrophils were isolated from murine bone marrow and conditioned media from stromal cells were generated as described above. Next, 200 µl of conditioned medium was added to the cancer cells containing neutrophils at the indicated ratios. After 24 hours of co-culture, the media were removed and the EO771/luc^+^ cells were washed with PBS. Subsequently, PBS containing luciferin (Promega, 150 µg/ml) was added to the wells. After 5 min of incubation, the chemiluminescence intensity was measured (IVIS Spectrum In-vivo-imaging system). The percent decrease in chemiluminescence intensity compared to EO771/luc^+^ cells without neutrophils reflects the cytotoxicity of neutrophils.

### 4. Flow cytometry

Cells (10^6^) were incubated with 100 μL staining mix containing 1 μL live-dead dye (Tag-it violet) in 1 ml D-PBS with 10^7^ cells. After incubation for 20 min at 37°C, the cells were washed twice with 10 ml D-PBS and centrifuged at 500g for 5 min each. Staining with various antibodies was performed in FACS buffer containing D-PBS, 5% FCS, and penicillin/streptomycin for 30 min at 4°C. The cells were washed with 200 μl FACS buffer and centrifuged at 500g for 5 min. Cells were resuspended in 100 μL FACS buffer. Unstained cells were used as negative controls. Occasionally, biotin labeled antibodies were used. For this, a second round with the secondary antibody was performed in a manner similar to the staining with labeled antibodies. Analysis was performed using the FlowJo software (RRID:SCR_008520, BD Biosciences).

### 5. Cell sorting

Stromal subpopulations were sorted from flushed bone marrow that was subjected to ACK-lysis in order to remove the red blood cells and from which CD45 cells were depleted using beads as described above. After washing, a mixture of the antibodies to be used including CD45 in addition to a live-dead stain were added for 30 minutes at 4°C. The cells were always sorted for high purity and collected in αMEM+10%FCS.

To assess apoptosis of B16 cells, staining for the annexin V-propidium iodide staining was performed using annexin-v conjugated to Alexa 647 (#640912 Biolegend, 1:50) in buffer containing 10mM HEPES pH 7.4, 140mM NaCl and 2.5mM CaCl_2_ for 20 minutes. This was stopped with an equal volume of the buffer and followed by the addition of propidium iodide 5 minutes before measurement of the corresponding sample. To evaluate cells proliferation, cells were fixed in 1% paraformaldehyde and permeabilized with 0.1% triton X (Sigma-Aldrich). The cells were then blocked in 5% BSA (Roth) for 15 minutes before staining them with an antibody directed against Ki67 (#652404, Biolegend, RRID:AB_2561525) for 30 minutes. The following anti-mouse antibodies and dyes were used:

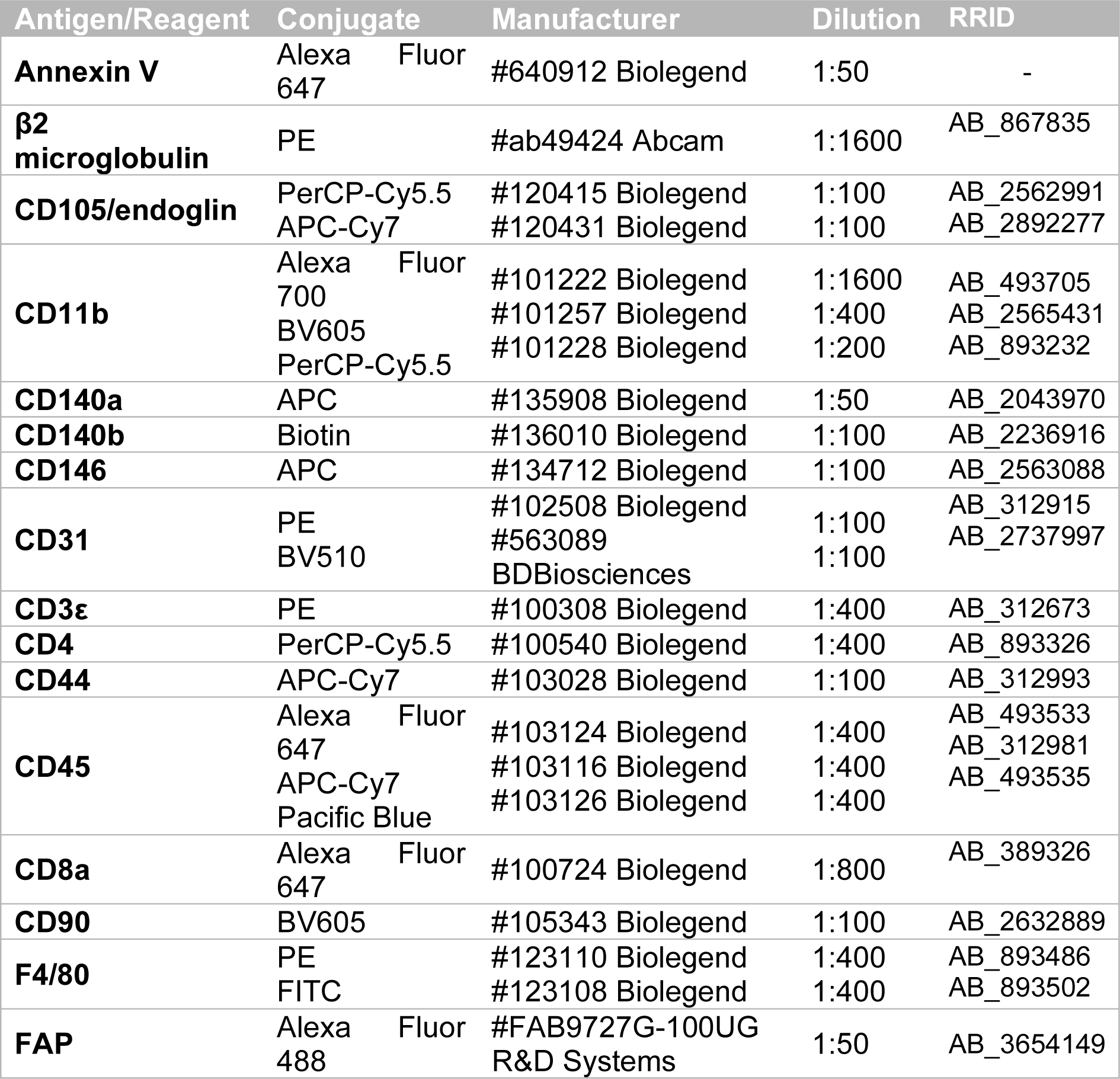
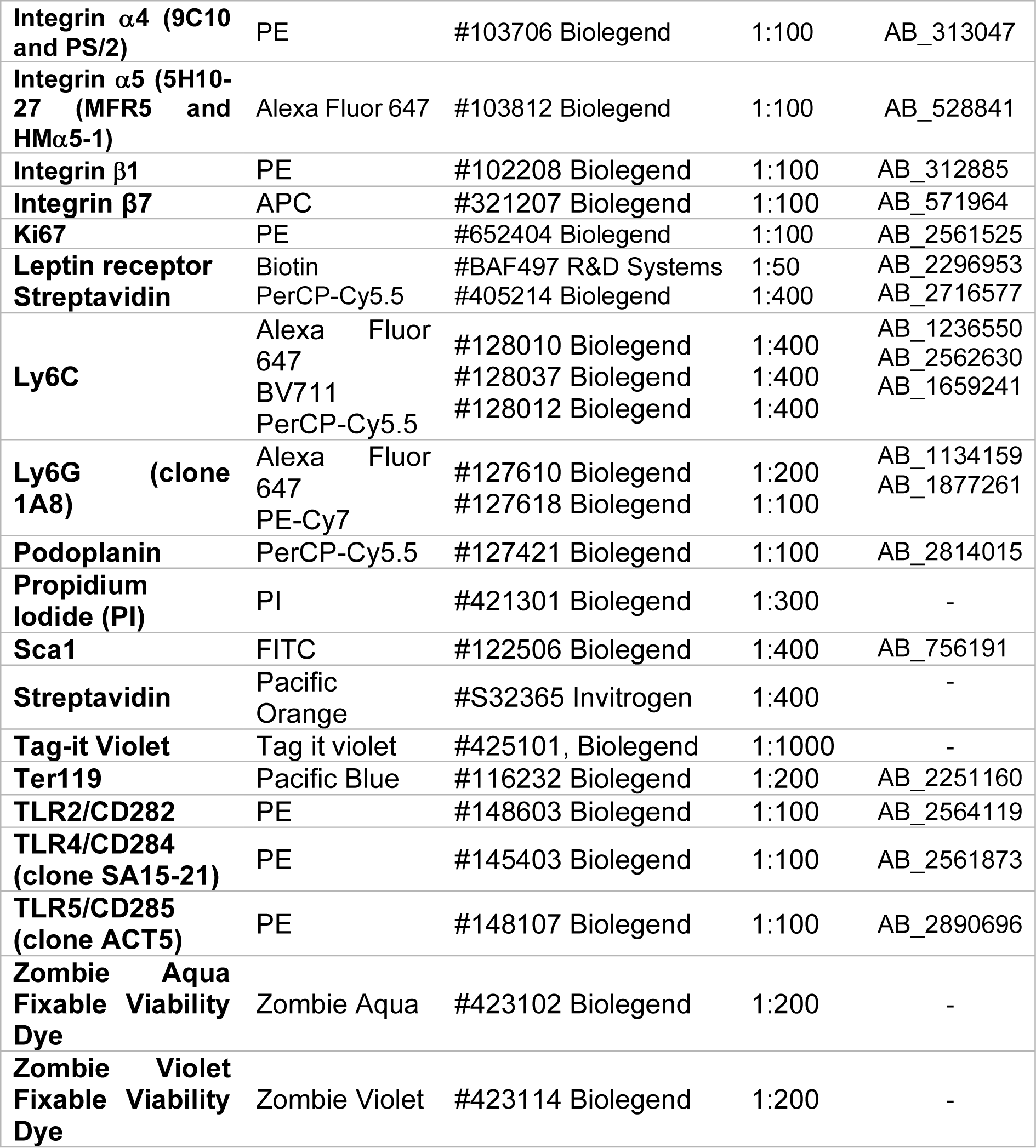

### 6. RNA Analysis

RNA was isolated using RNAzol (Sigma Aldrich) and reverse transcribed with a protocol using oligo(dT) primers (25 ng/μl), dNTPS (10 mM), RevertAid Reverse Transcriptase (200 U/μL, Thermo Fisher), and RiboLock RNase Inhibitor (40 U/μL, Thermo Fisher). DNA was digested for some primer pairs using an RNase-free DNase Set (79256, Qiagen). Subsequently, qPCR was performed using SensiFast Probe No-ROX (Bioline). qPCR results were normalized to those of murine HPRT. Where available, the probes and primers suggested by the universal probe library were used. The primers and probes used were as follows:

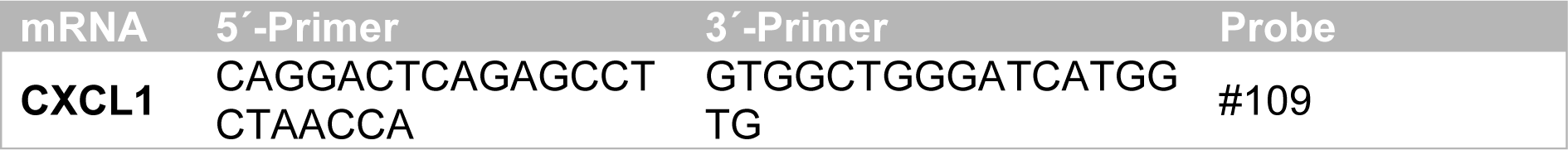
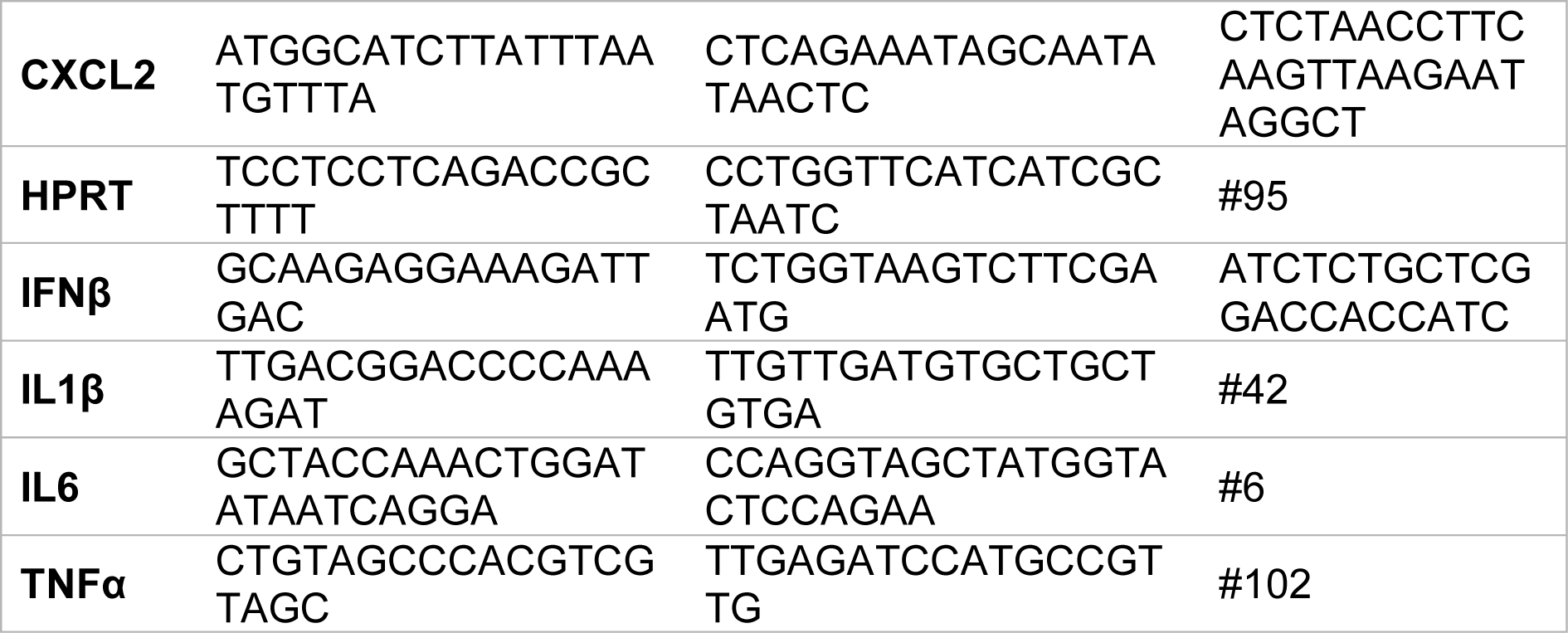

### 7. Protein analysis

#### Western blotting

Stromal cells were seeded in αMEM containing 10% FCS and 1% penicillin/streptomycin for 24 hours. The cells were then serum-starved overnight and treated with plasma fibronectin and the CS1 peptide at the indicated concentrations and time points. To assess NF-κB translocation, stromal cells were fractionated into cytoplasmic and nuclear fractions. For whole lysates, GAPDH was used as a loading control, whereas Histone H3 was used in the nuclear extracts. SDS-PAGE (10%) was performed and the following proteins were evaluated:

| Antibody | Clone | Manufacturer | RRID |
| --- | --- | --- | --- |
| **AKT** | Polyclonal | #9272 Cell Signaling | AB_329827 |
| **ERK 1/2** | Polyclonal | #9102 Cell Signaling | AB_330744 |
| **FAK** | Polyclonal | #3285 Cell Signaling | AB_2269034 |
| **GAPDH** | Polyclonal | #G9545 Sigma-Aldrich | AB_796208 |
| **Histone H3** | Polyclonal | #9715 Cell Signaling | AB_331563 |
| **IκBα** | 44D4 | #4812 Cell Signaling | AB_10694416 |
| **Integrin β1** | MB1.2 | #MAB1997 Millipore | AB_2128202 |
| **JNK/SAPK** | Polyclonal | #9252 Cell Signaling | AB_2250373 |
| **NF-κB (p65)** | C22B4 | #4764 Cell Signaling | AB_823578 |
| **phos- IκBα (Ser32)** | 14D4 | #2859 Cell Signaling | AB_561111 |
| **phos-AKT (Ser473)** | Polyclonal | #9271 Cell Signaling | AB_329825 |
| **phos-ERK 1/2 (Thr 202/Tyr 204)** | Polyclonal | #4376 Cell Signaling | AB_331772 |
| **phos-FAK (Tyr 397)** | D20B1 | #8556 Cell Signaling | AB_10891442 |
| **phos-JNK (Thr 183/Tyr 185)** | Polyclonal | #9251 Cell Signaling | AB_331659 |
| **phos-NF-κB (Ser536)** | 93H1 | #3033 Cell Signaling | AB_331284 |

The secondary antibodies used were: goat anti-rabbit (HRP, 111-035-045, RRID: AB\_2337938) or goat anti-rat (HRP, 112-036-071, RRID: AB\_2338144)(Jackson ImmunoResearch). The application of ECL western blotting-substrate was followed by detection of the chemiluminescence signal with Fusion FX7 (Vilber).

#### Fibronectin ELISA

The stromal cells were isolated and seeded in FCS-free αMEM for 24 hours. The media were collected and protein content, used for adjustment of values, was measured using BCA (Pierce). Plates were coated with the primary antibody against fibronectin (F3648, Sigma-Aldrich, polyclonal, 0.12 μg/ml, RRID: AB_476976). The standard used was murine plasma (#IFMBN, Dunn). The secondary anti-fibronectin antibody was conjugated to HRP (#IRBAMSFBNGFHRP1MG, Loxo, RRID: AB_11043433) as published^20,48^.

#### Proteomic evaluation

Osterix-expressing cells were sorted based on tdTomato expression in cre-expressing cells and the remaining cells were used as controls. Cell pellets were lysed and digested. LC-MS/MS measurements were performed using an Easy nLC 1200 (Thermo Scientific) coupled to a timsTOF Pro (Bruker Daltonics). The mass spectrometer was operated in the data-dependent PASEF mode. Raw data were processed using the MaxQuant computational platform version 2.2.0.0 (RRID:SCR_014485). Label-free quantification (LFQ) intensities were obtained for each protein. Data were analyzed using the STRING DB platform (RRID:SCR_005223).

### 8. Statistical analysis

Statistical analyses were performed using GraphPad Prism version 8 (RRID:SCR_002798). Analysis of variance (ANOVA) was used when to compare more than two groups as appropriate. If the global probability was less than 5%, comparisons between pairs were performed, as appropriate. Most data were compared using unpaired *t*-tests, except for qPCR data involving the expression of cytokines, where the Mann-Whitney-test was used. Analysis of the transwell assay data was performed using one-way ANOVA in conjunction with a Bonferroni post-test.

## Ethics statement

All animal protocols were approved as outlined above. No patient data are included.

## CRediT authorship contribution statement

**Alexander Lubosch:** Writing – review and editing, Visualization, Methodology, Investigation, conceptualization. **Lauren Pitt:** Writing – review and editing, Visualization, Methodology, Investigation. **Caren Zoeller:** Writing – review and editing, Visualization, Methodology, Investigation. **Franziska Wirth:** Writing – review and editing, Visualization, Validation, Investigation. **Tarik Exner:** Writing – review and editing, Visualization, Methodology. **Barbara Steigenberger:** Writing – review and editing, Visualization, Methodology, Investigation. **Jutta Schroeder-Braunstein:** Writing – review & editing, Visualization. **Guido Wabnitz:** Writing – review and editing, Methodology, Investigation. **Inaam A. Nakchbandi:** Writing – review and editing, Writing – original draft, Visualization, Supervision, Resources, Project administration, Methodology, Funding acquisition, Conceptualization.

## Data availability statement

The data generated in this study are available within the article and its supplementary data files.

Data from proteomic analysis are presented in the supplementary material. Raw data are available upon reasonable request by e-mail from the corresponding author. Please write in the subject line: “Request for raw data-2024”, and specify which data are needed in the e-mail you send. Please allow for two weeks to receive a response.

## Supplemental information

One document with 6 Supplementary Figures and 1 Supplementary Table.

*Supplementary Figure 1.*

*A-B. Characteristics of depleted stromal cells.*

*C-D. Tumor volume decreases in the presence of stromal cells in two models (weight data are shown in Figure 1).*

*E-F. Absence of changes in CD45+ and T-cell numbers in the tumors.*

*G-H. Tumor volume decreases in the absence of mature T-cells (weight data are shown in Figure 1).*

*I. Amount of stromal cells detected after 3 days in vivo.*

*Supplementary Figure 2.*

*A-B. Inhibition of growth is specific to bone marrow stromal cells*

*C. Sorting of cancer associated fibroblasts (CAFs)*

*Supplementary Figure 3.*

*Heat map and volcano plot for differentially expressed molecules between osterix expressing cells and those not expressing osterix.*

*Supplementary Figure 4.*

*Effect of CS1 on B16 tumor cells and stromal cells.*

*Supplementary Figure 5.*

*A. Immune cell changes in the presence of stromal cells*

*B-E. CS1 enhances Ly6G+ cell migration and affects activity in vitro*

*Supplementary Figure 6.*

*A-B. CS1 increases NF-κB signaling*

*C. Expression of some of the receptors known to mediate fibronectin effects on cells and to interact with NF-κB.*

*Supplementary Table 1.*

*List of differentially expressed molecules between osterix expressing cells and those not expressing osterix*

## Acknowledgements

We thank Prof. Reinhard Fässler for invaluable support, and Prof. Axel Roers for scientific input. We acknowledge funding from the German Research Council (DFG) (NA-400/9; NA-400/10-401246035) and the Max-Planck Society (M.KF.A.BIOC0001/K440).

## References

1. Ping, Q., Yan, R., Cheng, X., Wang, W., Zhong, Y., Hou, Z., Shi, Y., Wang, C., and Li, R. (2021). Cancer-associated fibroblasts: overview, progress, challenges, and directions. Cancer Gene Ther 28, 984–999. 10.1038/s41417-021-00318-4.

2. Bu, L., Baba, H., Yoshida, N., Miyake, K., Yasuda, T., Uchihara, T., Tan, P., and Ishimoto, T. (2019). Biological heterogeneity and versatility of cancer-associated fibroblasts in the tumor microenvironment. Oncogene 38, 4887–4901. 10.1038/s41388-019-0765-y.

3. Carpenter, E.S., Vendramini-Costa, D.B., Hasselluhn, M.C., Maitra, A., Olive, K.P., Cukierman, E., Pasca di Magliano, M., and Sherman, M.H. (2024). Pancreatic Cancer-Associated Fibroblasts: Where Do We Go from Here? Cancer Res 84, 3505–3508. 10.1158/0008-5472.CAN-24-2860.

4. Hernandez, J.L., Padilla, L., Dakhel, S., Coll, T., Hervas, R., Adan, J., Masa, M., Mitjans, F., Martinez, J.M., Coma, S., et al. (2013). Therapeutic targeting of tumor growth and angiogenesis with a novel anti-S100A4 monoclonal antibody. PLoS One 8, e72480. 10.1371/journal.pone.0072480.

5. Teichgraber, V., Monasterio, C., Chaitanya, K., Boger, R., Gordon, K., Dieterle, T., Jager, D., and Bauer, S. (2015). Specific inhibition of fibroblast activation protein (FAP)-alpha prevents tumor progression in vitro. Adv Med Sci 60, 264–272. 10.1016/j.advms.2015.04.006.

6. Özdemir, B.C., Pentcheva-Hoang, T., Carstens, J.L., Zheng, X., Wu, C.C., Simpson, T.R., Laklai, H., Sugimoto, H., Kahlert, C., Novitskiy, S.V., et al. (2014). Depletion of Carcinoma-Associated Fibroblasts and Fibrosis Induces Immunosuppression and Accelerates Pancreas Cancer with Reduced Survival. Cancer Cell 25, 719–734. 10.1016/J.CCR.2014.04.005.

7. Bhattacharjee, S., Hamberger, F., Ravichandra, A., Miller, M., Nair, A., Affo, S., Filliol, A., Chin, L.K., Savage, T.M., Yin, D., et al. (2021). Tumor restriction by type I collagen opposes tumor-promoting effects of cancer-associated fibroblasts. The Journal of Clinical Investigation 131. 10.1172/JCI146987.

8. Zhang, R., Qi, F., Zhao, F., Li, G., Shao, S., Zhang, X., Yuan, L., and Feng, Y. (2019). Cancer-associated fibroblasts enhance tumor-associated macrophages enrichment and suppress NK cells function in colorectal cancer. Cell Death and Disease 10. 10.1038/s41419-019-1435-2.

9. Gok Yavuz, B., Gunaydin, G., Gedik, M.E., Kosemehmetoglu, K., Karakoc, D., Ozgur, F., and Guc, D. (2019). Cancer associated fibroblasts sculpt tumour microenvironment by recruiting monocytes and inducing immunosuppressive PD-1 + TAMs. Scientific reports 9. 10.1038/s41598-019-39553-z.

10. Hutton, C., Heider, F., Blanco-Gomez, A., Banyard, A., Kononov, A., Zhang, X., Karim, S., Paulus-Hock, V., Watt, D., Steele, N., et al. (2021). Single-cell analysis defines a pancreatic fibroblast lineage that supports anti-tumor immunity. Cancer Cell 39, 1227–1244.e1220. 10.1016/J.CCELL.2021.06.017.

11. McFarlane, A.J., Fercoq, F., Coffelt, S.B., and Carlin, L.M. (2021). Neutrophil dynamics in the tumor microenvironment. The Journal of Clinical Investigation 131. 10.1172/JCI143759.

12. Kuwabara, W.M.T., Andrade-Silva, J., Pereira, J.N.B., Scialfa, J.H., and Cipolla-Neto, J. (2019). Neutrophil activation causes tumor regression in Walker 256 tumor-bearing rats. Scientific reports 9, 16524. 10.1038/s41598-019-52956-2.

13. Hinshaw, D.C., and Shevde, L.A. (2019). The Tumor Microenvironment Innately Modulates Cancer Progression. Cancer Res 79, 4557–4566. 10.1158/0008-5472.CAN-18-3962.

14. Yang, D., Liu, J., Qian, H., and Zhuang, Q. (2023). Cancer-associated fibroblasts: from basic science to anticancer therapy. Exp Mol Med 55, 1322–1332. 10.1038/s12276-023-01013-0.

15. Ricci, B., Tycksen, E., Celik, H., Belle, J.I., Fontana, F., Civitelli, R., and Faccio, R. (2020). Osterix-Cre marks distinct subsets of CD45- and CD45+stromal populations in extra-skeletal tumors with pro-tumorigenic characteristics. Elife 9. 10.7554/eLife.54659.

16. Rossnagl, S., Ghura, H., Groth, C., Altrock, E., Jakob, F., Schott, S., Wimberger, P., Link, T., Kuhlmann, J.D., Stenzl, A., et al. (2018). A Subpopulation of Stromal Cells Controls Cancer Cell Homing to the Bone Marrow. Cancer Res 78, 129–142. 10.1158/0008-5472.CAN-16-3507.

17. Segre, J.A., Nemhauser, J.L., Taylor, B.A., Nadeau, J.H., and Lander, E.S. (1995). Positional cloning of the nude locus: genetic, physical, and transcription maps of the region and mutations in the mouse and rat. Genomics 28, 549–559. S0888-7543(85)71187-1 [pii]. 10.1006/geno.1995.1187.

18. Ghura, H., Keimer, M., von Au, A., Hackl, N., Klemis, V., and Nakchbandi, I.A. (2021). Inhibition of fibronectin accumulation suppresses tumor growth. Neoplasia 23, 837–850. 10.1016/j.neo.2021.06.012.

19. Rossnagl, S., Altrock, E., Sens, C., Kraft, S., Rau, K., Milsom, M.D., Giese, T., Samstag, Y., and Nakchbandi, I.A. (2016). EDA-Fibronectin Originating from Osteoblasts Inhibits the Immune Response against Cancer. PLoS Biol 14, e1002562. 10.1371/journal.pbio.1002562.

20. von Au, A., Vasel, M., Kraft, S., Sens, C., Hackl, N., Marx, A., Stroebel, P., Hennenlotter, J., Todenhofer, T., Stenzl, A., et al. (2013). Circulating fibronectin controls tumor growth. Neoplasia 15, 925–938.

21. Yoshida, S., Asanoma, K., Yagi, H., Onoyama, I., Hori, E., Matsumura, Y., Okugawa, K., Yahata, H., and Kato, K. (2021). Fibronectin mediates activation of stromal fibroblasts by SPARC in endometrial cancer cells. BMC Cancer 21, 156. 10.1186/s12885-021-07875-9.

22. Wirth, F., Zoeller, C., Lubosch, A., Schroeder-Braunstein, J., Wabnitz, G., and Nakchbandi, I.A. (2024). Insights into the metastatic bone marrow niche gained from fibronectin and beta1 integrin transgenic mice. Neoplasia 58, 101058. 10.1016/j.neo.2024.101058.

23. Wirth, F., Huck, K., Lubosch, A., Zoeller, C., Ghura, H., Porubsky, S., and Nakchbandi, I.A. (2021). Cdc42 in osterix-expressing cells alters osteoblast behavior and myeloid lineage commitment. Bone 153, 116150–116150. 10.1016/j.bone.2021.116150.

24. Klemis, V., Ghura, H., Federico, G., Wurfel, C., Bentmann, A., Gretz, N., Miyazaki, T., Grone, H.J., and Nakchbandi, I.A. (2017). Circulating fibronectin contributes to mesangial expansion in a murine model of type 1 diabetes. Kidney Int 91, 1374–1385. 10.1016/j.kint.2016.12.006.

25. Bentmann, A., Kawelke, N., Moss, D., Zentgraf, H., Bala, Y., Berger, I., Gasser, J.A., and Nakchbandi, I.A. (2010). Circulating fibronectin affects bone matrix, whereas osteoblast fibronectin modulates osteoblast function. J Bone Miner Res 25, 706–715. 10.1359/jbmr.091011.

26. Hofbauer, L.C., Bozec, A., Rauner, M., Jakob, F., Perner, S., and Pantel, K. (2021). Novel approaches to target the microenvironment of bone metastasis. Nat Rev Clin Oncol 18, 488–505. 10.1038/s41571-021-00499-9.

27. Leiss, M., Beckmann, K., Giros, A., Costell, M., and Fassler, R. (2008). The role of integrin binding sites in fibronectin matrix assembly in vivo. Current opinion in cell biology 20, 502–507. S0955-0674(08)00111-7 [pii] 10.1016/j.ceb.2008.06.001.

28. Schumacher, S., Dedden, D., Nunez, R.V., Matoba, K., Takagi, J., Biertumpfel, C., and Mizuno, N. (2021). Structural insights into integrin alpha(5)beta(1) opening by fibronectin ligand. Sci Adv 7. 10.1126/sciadv.abe9716.

29. Kamarajan, P., Garcia-Pardo, A., D’Silva, N.J., and Kapila, Y.L. (2010). The CS1 segment of fibronectin is involved in human OSCC pathogenesis by mediating OSCC cell spreading, migration, and invasion. BMC Cancer 10, 330. 10.1186/1471-2407-10-330.

30. Sens, C., Altrock, E., Rau, K., Klemis, V., von Au, A., Pettera, S., Uebel, S., Damm, T., Tiwari, S., Moser, M., and Nakchbandi, I.A. (2017). An O-Glycosylation of Fibronectin Mediates Hepatic Osteodystrophy Through alpha4beta1 Integrin. J Bone Miner Res 32, 70–81. 10.1002/jbmr.2916.

31. Rose, D.M., Han, J., and Ginsberg, M.H. (2002). Alpha4 integrins and the immune response. Immunological reviews 186, 118–124. 10.1034/j.1600-065x.2002.18611.x.

32. Boivin, G., Faget, J., Ancey, P.B., Gkasti, A., Mussard, J., Engblom, C., Pfirschke, C., Contat, C., Pascual, J., Vazquez, J., et al. (2020). Durable and controlled depletion of neutrophils in mice. Nat Commun 11, 2762. 10.1038/s41467-020-16596-9.

33. Stegelmeier, A.A., Darzianiazizi, M., Hanada, K., Sharif, S., Wootton, S.K., Bridle, B.W., and Karimi, K. (2021). Type I Interferon-Mediated Regulation of Antiviral Capabilities of Neutrophils. Int J Mol Sci 22. 10.3390/ijms22094726.

34. Sens, C., Huck, K., Pettera, S., Uebel, S., Wabnitz, G., Moser, M., and Nakchbandi, I.A. (2017). Fibronectins containing extradomain A or B enhance osteoblast differentiation via distinct integrins. J Biol Chem 292, 7745–7760. 10.1074/jbc.M116.739987.

35. Ambesi, A., Maddali, P., and McKeown-Longo, P.J. (2022). Fibronectin Functions as a Selective Agonist for Distinct Toll-like Receptors in Triple-Negative Breast Cancer. Cells 11, 2074–2074. 10.3390/cells11132074.

36. Wirth, F., Lubosch, A., Hamelmann, S., and Nakchbandi, I.A. (2020). Fibronectin and Its Receptors in Hematopoiesis. Cells 9, 2717–2717. 10.3390/cells9122717.

37. Selders, G.S., Fetz, A.E., Radic, M.Z., and Bowlin, G.L. (2017). An overview of the role of neutrophils in innate immunity, inflammation and host-biomaterial integration. Regen Biomater 4, 55–68. 10.1093/rb/rbw041.

38. Deligne, C., Murdamoothoo, D., Gammage, A.N., Gschwandtner, M., Erne, W., Loustau, T., Marzeda, A.M., Carapito, R., Paul, N., Velazquez-Quesada, I., et al. (2020). Matrix-Targeting Immunotherapy Controls Tumor Growth and Spread by Switching Macrophage Phenotype. Cancer Immunol Res 8, 368–382. 10.1158/2326-6066.CIR-19-0276.

39. Gao, H.X., Bhattacharya, S., Matheny, C.J., Yanamandra, N., Zhang, S.Y., Emerich, H., Li, Y.F., Bojczuk, P., Shi, H., Wang, W., et al. (2018). Synergy of TLR4 agonist GSK1795091, an innate immune activator, with agonistic antibody against co-stimulatory immune checkpoint molecule OX40 in cancer immunotherapy. Journal of Clinical Oncology 36. DOI 10.1200/JCO.2018.36.15_suppl.12055.

40. Oblak, A., and Jerala, R. (2011). Toll-like receptor 4 activation in cancer progression and therapy. Clinical & developmental immunology 2011, 609579. 10.1155/2011/609579.

41. Luchner, M., Reinke, S., and Milicic, A. (2021). TLR Agonists as Vaccine Adjuvants Targeting Cancer and Infectious Diseases. Pharmaceutics 13. 10.3390/pharmaceutics13020142.

42. Ollauri-Ibanez, C., Nunez-Gomez, E., Egido-Turrion, C., Silva-Sousa, L., Diaz-Rodriguez, E., Rodriguez-Barbero, A., Lopez-Novoa, J.M., and Pericacho, M. (2020). Continuous endoglin (CD105) overexpression disrupts angiogenesis and facilitates tumor cell metastasis. Angiogenesis 23, 231–247. 10.1007/s10456-019-09703-y.

43. Gershkovitz, M., Caspi, Y., Fainsod-Levi, T., Katz, B., Michaeli, J., Khawaled, S., Lev, S., Polyansky, L., Shaul, M.E., Sionov, R.V., et al. (2018). TRPM2 mediates neutrophil killing of disseminated tumor cells. Cancer Research 78, 2680–2690. 10.1158/0008-5472.CAN-17-3614/653280/AM/TRPM2-MEDIATES-NEUTROPHIL-KILLING-OF-DISSEMINATED.

44. Chung, J.Y., Tang, P.C., Chan, M.K., Xue, V.W., Huang, X.R., Ng, C.S., Zhang, D., Leung, K.T., Wong, C.K., Lee, T.L., et al. (2023). Smad3 is essential for polarization of tumor-associated neutrophils in non-small cell lung carcinoma. Nat Commun 14, 1794. 10.1038/s41467-023-37515-8.

45. Mishalian, I., Bayuh, R., Levy, L., Zolotarov, L., Michaeli, J., and Fridlender, Z.G. (2013). Tumor-associated neutrophils (TAN) develop pro-tumorigenic properties during tumor progression. Cancer Immunology, Immunotherapy 62, 1745–1756. 10.1007/S00262-013-1476-9/FIGURES/5.

46. Rossnagl, S., von Au, A., Vasel, M., Cecchini, A.G., and Nakchbandi, I.A. (2014). Blood clot formation does not affect metastasis formation or tumor growth in a murine model of breast cancer. PLoS One 9, e94922. 10.1371/journal.pone.0094922.

47. Kawelke, N., Vasel, M., Sens, C., von Au, A., Dooley, S., and Nakchbandi, I.A. (2011). Fibronectin Protects from Excessive Liver Fibrosis by Modulating the Availability of and Responsiveness of Stellate Cells to Active TGF-beta. PLoS One 6, e28181. 10.1371/journal.pone.0028181.PONE-D-11-09211 [pii].

48. Hackl, N.J., Bersch, C., Feick, P., Antoni, C., Franke, A., Singer, M.V., and Nakchbandi, I.A. (2010). Circulating fibronectin isoforms predict the degree of fibrosis in chronic hepatitis C. Scand J Gastroenterol 45, 349–356. 10.3109/00365520903490606.

