## Supplementary file (Figures and Table) for "The Immune Response Against Cancer is Modulated by Stromal Cell Fibronectin"

### Supplementary Figure 1

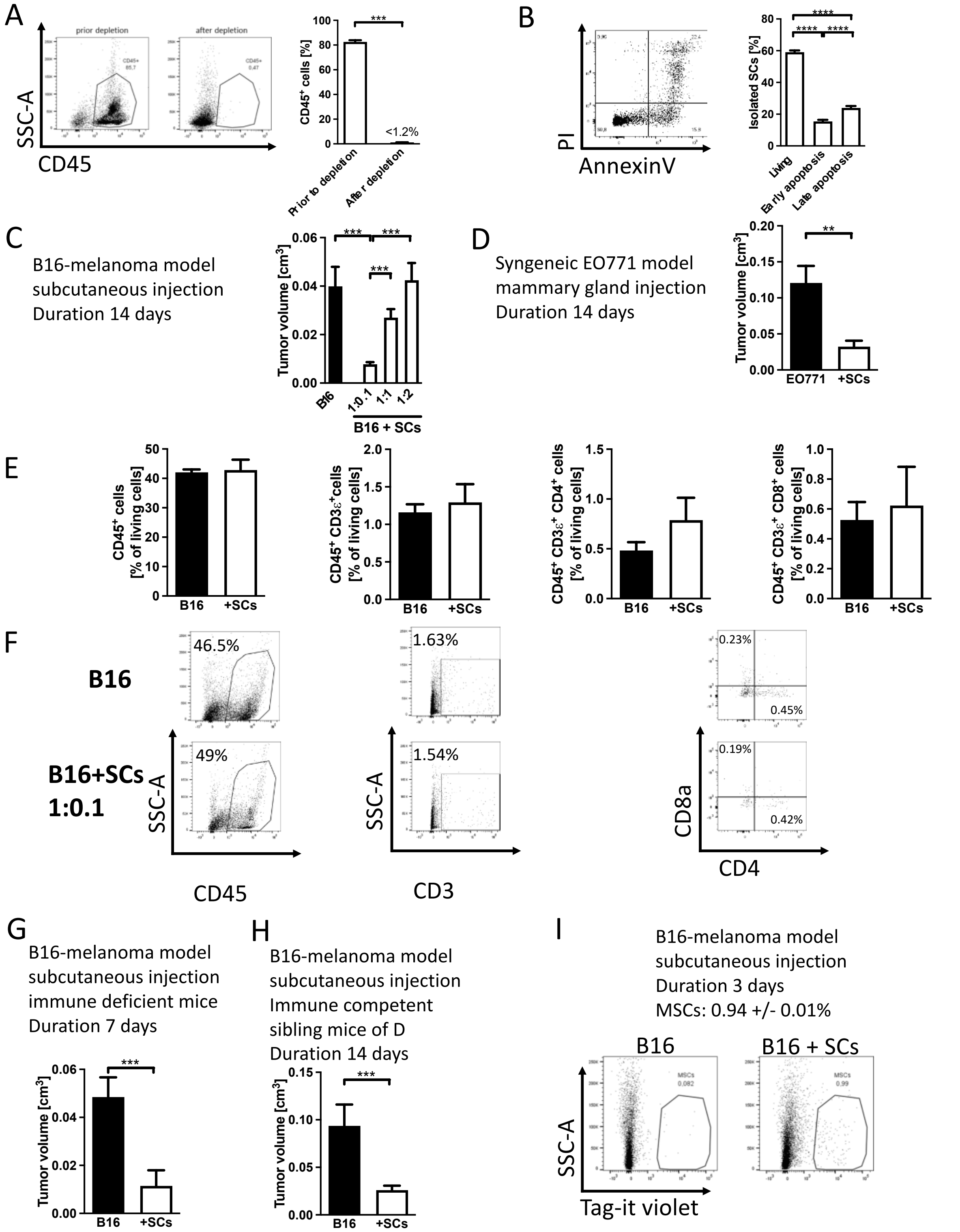


***Supplementary Figure 1.***
***A-B. Characteristics of depleted stromal cells.***

A. Flow cytometry shows that bone marrow cells depleted of $CD45^+$ immune cells contains less than 1% of $CD45^+$ cells. N=20/20. B. Viability after depletion is shown. N=13/13/13. ****p<0.0001.

Methods:
Bone marrow cells were flushed using D-PBS. This was followed by depletion of $CD45^+$ cells using magnetic beads precoated with labeled CD45 antibody. Evaluation of stromal cells after depletion was performed by staining the stromal cells with a labeled CD45 antibody and parallel samples with an appropriate secondary antibody (Mouse anti-rat labeled with FITC, Biolegend #407505). To determine apoptosis, the cells were stained against annexin-v antibody labeled with alexa 647 (#640912 Biolegend, 1:50) in Annexin-binding-buffer containing 10mM HEPES pH 7.4, 140mM NaCl and 2.5mM $CaCl_2$ for 20 minutes before stopping the reaction with an equal volume of binding buffer. Five minutes before analysis, propidium iodide was added to the cells.

***C-D. Tumor volume decreases in the presence of stromal cells in two models (weight data are shown in Figure 1).***
Schematic of the treatment regimen I shown in Figure 1A. Bone marrow cells were isolated, depleted of immune cells, and mixed in different ratios with tumor cells before injecting the mixture subcutaneously. Growth was evaluated after 14 days. C. Injection of stromal cells with B16 melanoma tumor cells results in smaller tumors when the ratio is 1:0.1 (Tumor cells:stromal cells). While The ratio of 1:1 still led to a small decrease in tumor volume, the 1:2 ratio failed to inhibit cancer growth. N=12/12/12/12, *p<0.05, ***p<0.001. D. Using syngeneic breast tumor cells (EO771) at a ratio of 1:0.1 injected orthotopically in the mammary gland confirms the decrease in growth. N=10/8, **p<0.01, ***p<0.001.

***E-F. Absence of changes in $CD45^+$ and T-cell numbers in the tumors.***
C. Stromal cells (SC) were mixed with B16 cancer cells at a ratio of 1:0.1 (B16/SCs). After 14 days, tumors were removed, digested, cells stained for the markers shown and evaluated by flow cytometry. No difference in total number of immune cells ($CD45^+$) or in the various T-cell markers in the experimental groups could be detected. $CD45^+$: N=12/12, $CD45^+CD3\varepsilon^+$: N=12/6, $CD45^+CD3\varepsilon^+CD4^+$: N=12/6, $CD45^+CD3\varepsilon^+CD8^+$: N=12/6. D. Representative pictures of flow cytometry evaluation.

The antibodies used were:
CD3ε (PE, #100308 Biolegend; dilution 1:400)
CD4 (PerCP-Cy5.5, #100540 Biolegend; dilution 1:400)
CD8a (Alexa Fluor 647, #100724 Biolegend; dilution 1:800)

***G-H. Tumor volume decreases in the absence of mature T-cells (weight data are shown in Figure 1).***
Schematic of the treatment regimen I shown in Figure 1A. Bone marrow cells were isolated, depleted of immune cells, and mixed in different ratios with tumor cells before injecting the mixture subcutaneously. Growth was evaluated after 7-14 days. G. Tumor in the presence of stromal cells is maintained using B16 melanoma cells in animals lacking T cells on a different genetic background. N=8/8, **p<0.01, ***p<0.001. H. The genetic background of the mice presented in D did not prevent the inhibitory effect of stromal cells. N=9/15, **p<0.01.

***I. Amount of stromal cells detected after 3 days in vivo.***
Stromal cells were labeled before mixing with B16 tumor cells and injection subcutaneously. Three days after injection only 1% of the cells in the tumor were labeled suggesting that few cells are sufficient for the inhibitory effect. N=4/5, *p<0.05.

### Supplementary Figure 2

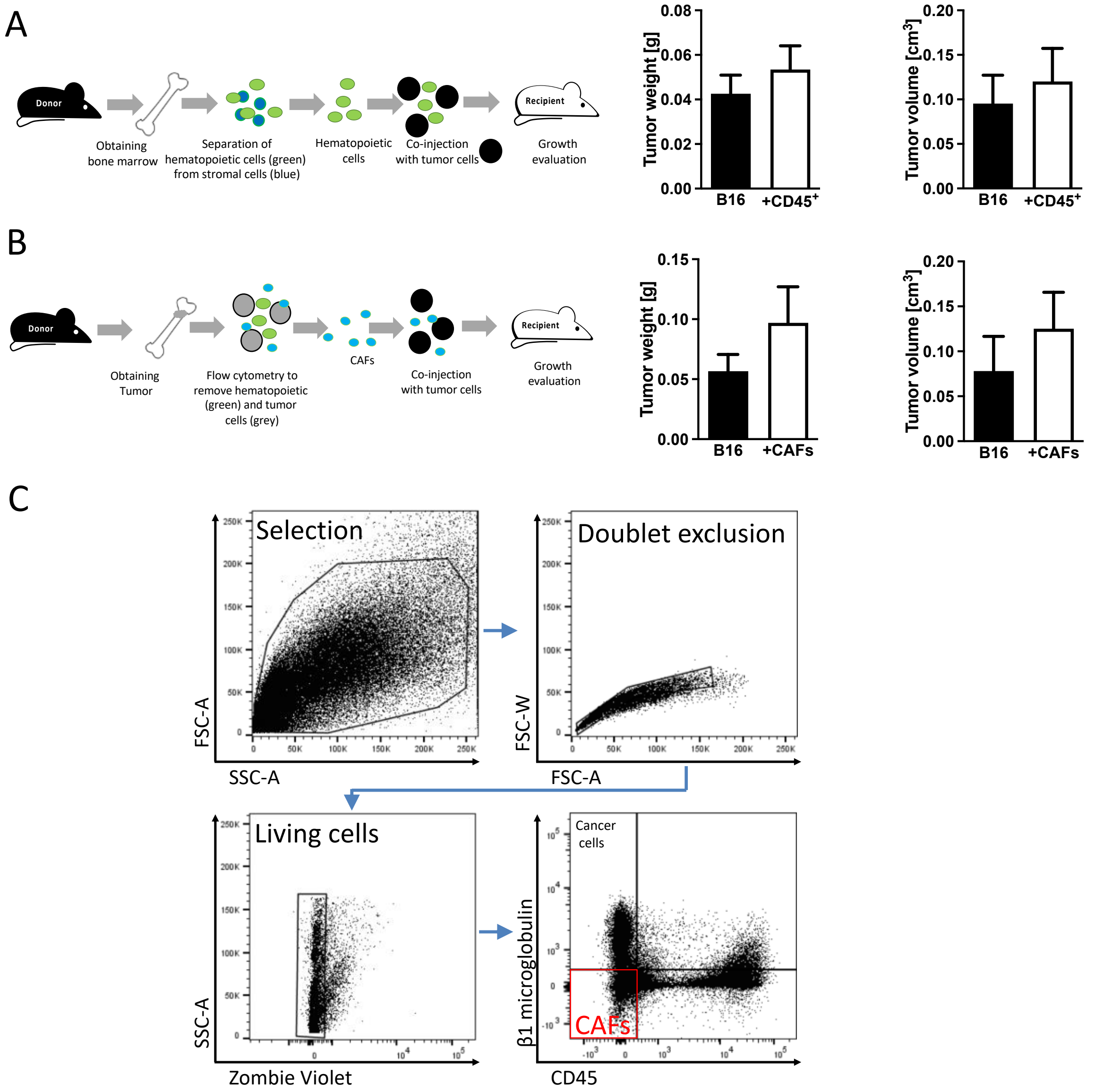


***Supplementary Figure 2.***

***A-B. Inhibition of growth is specific to bone marrow stromal cells***

A. Using $CD45^+$ cells instead of stromal cells did not result in growth inhibition. N=5/10. B. Cancer-associated fibroblasts (CAFs) did not affect a decrease or an increase in cancer growth in our model. CAFs were isolated from tumors of human breast cancer injected into the bone marrow of mice to induce bone metastases. Tumors were digested, stained, and sorted by excluding immune cells ($CD45^+$) and the human tumor cells (by using a human specific antibody). CAFs were mixed with B16 tumor cells at a ratio of 1:0.1 (B16:CAFs) before injecting them into mice subcutaneously. N=13/15. Comparisons using t-tests were performed.

***C. Sorting of cancer associated fibroblasts (CAFs)***

Human breast cancer cells were injected into the bone marrow to induce bone metastatic lesions. After 6 weeks, the tumors were isolated, digested, stained, and CAFs were sorted by excluding immune cells ($CD45^+$) and the human cancer cells (by expression of the human specific β2-microglobulin). The gating strategy is shown.

Methods: **Isolation of cancer-associated fibroblasts (CAFs)**
CAFs were isolated from excess material of intratibial MDA-MB-231-B/luc$^+$ (RRID:CVCL_0062) tumors from another study (G-261/20) [49]. Murine immune cells were stained for CD45 while human MDA-MB-231-B/luc$^+$ cells were stained for β2 microglobulin (clone B2M-01, Abcam). Cells negative for both markers were then defined as CAFs and used for the experiments.

### Supplementary Figure 3

A

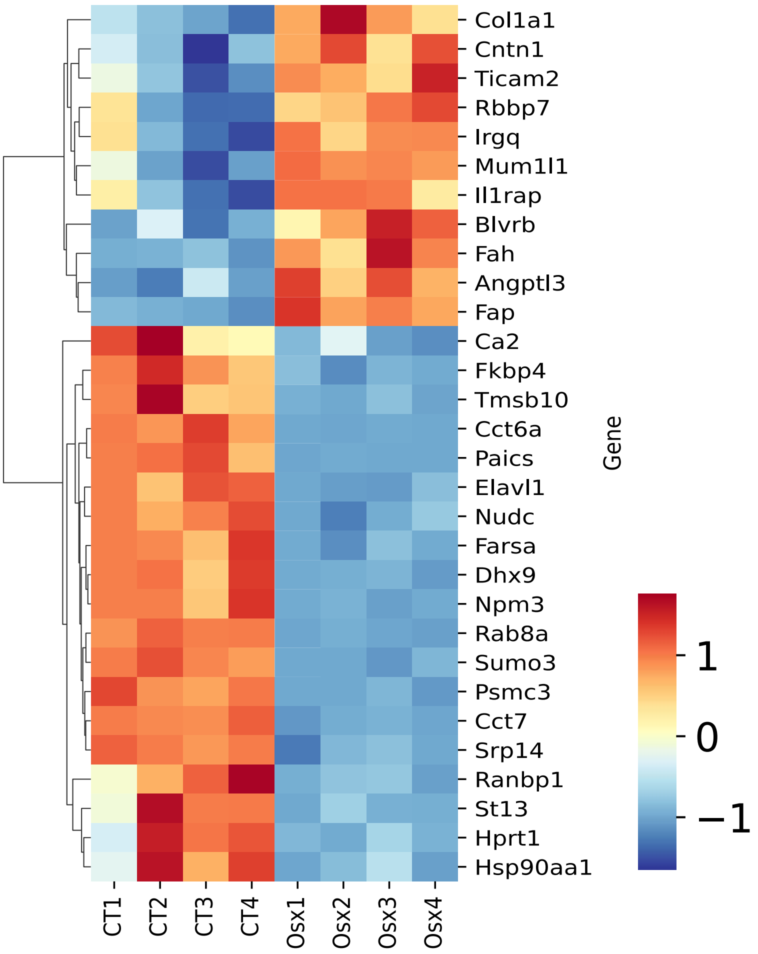


B

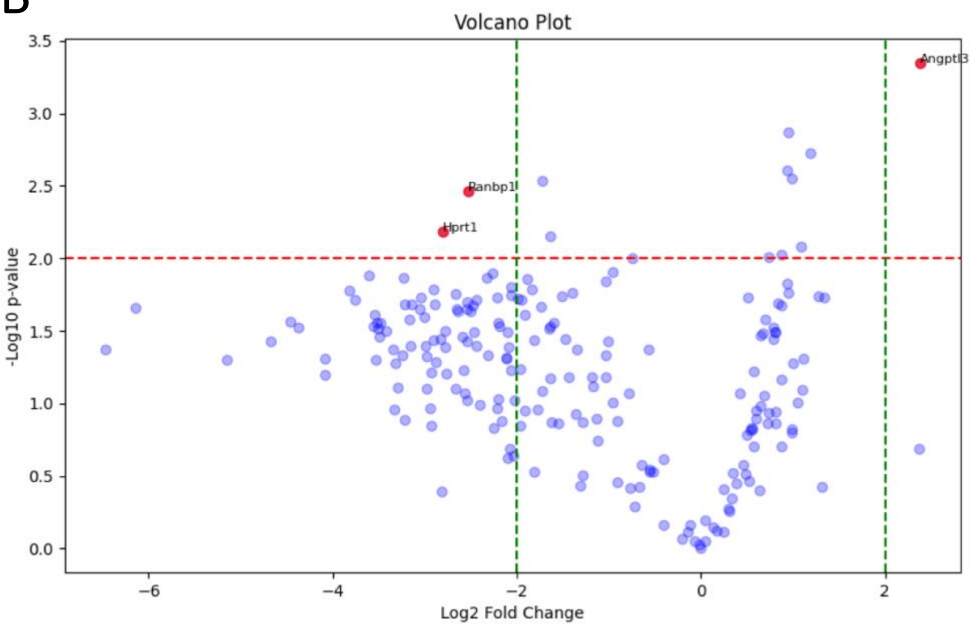

***Supplementary Figure 3. Heat map and volcano plot for differentially expressed molecules between osterix expressing cells and those not expressing osterix.*** Cells were isolated from the bone marrow of mice with the genotype osterix-cre_tdTomato$^{fl/+}$. Cells that express osterix-cre will remove the floxed stop codon at the beginning of the tdTomato transgene leading to tdTomato-stained cells that were detected by flow cytometry and sorted. These were compared to those that did not express tdTomato using proteomic analysis.(A) Heat map shows the results of evaluation of the normalized values. Since molecules with missing values were discarded in this analysis, the results differ some from Supplementary Table 1. (B) Molecules with $p<0.01$ and ratio difference of > 2 are shown in a volcano plot performed with raw data prior to normalization. N= 4/4.

**Additional information:**

The volcano plot (Supplementary Figure 3B) highlighted that three molecules were markedly different:

a. Ranbp1 (nuclear Ras binding protein-1) is a molecule involved in cell proliferation and is required for normal progression of mitosis (Li, Ng et al. 2007). Its decrease suggests less proliferation of cells that express osterix in comparison to those that do not.

b. HPRT1 (Hypoxanthine phosphoribosyltransferase 1) is required for purine nucleotide generation, in line with a role in proliferation and cell function (Vogel, Moehrle et al. 2019).

c. While Ranbp1 and HPRT1 were diminished, angptl3 (Angiopoietin like 3) was increased. A function attributed to this molecule is supporting proliferation (Camenisch, Pisabarro et al. 2002, Carbone, Piro et al. 2018).

These findings thus suggest that osterix-expressing stromal cells respond to proliferative cues differently from other stromal cells.

#### Supplementary Figure 4

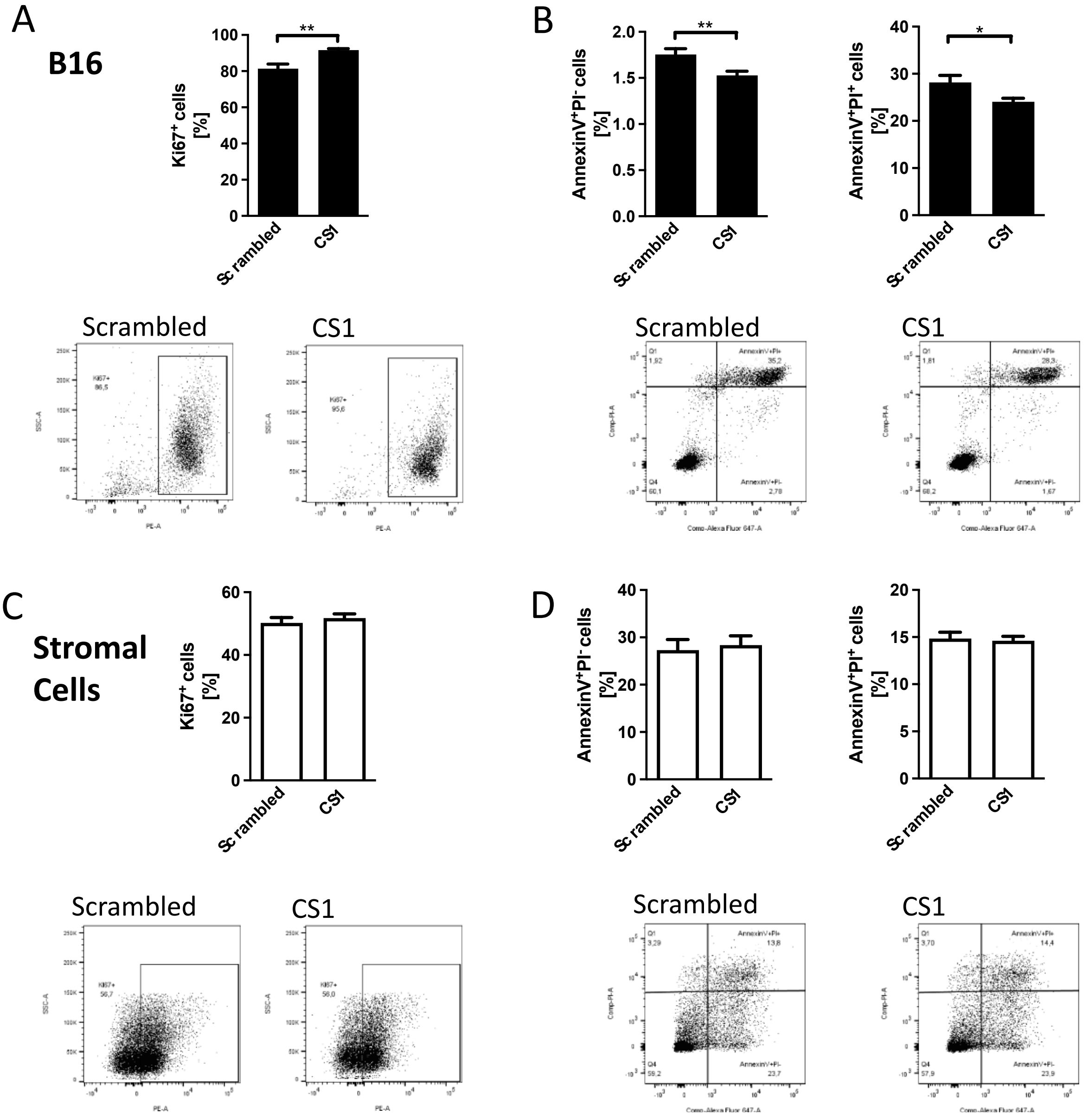


***Supplementary Figure 4. Effect of CS1 on B16 tumor cells and stromal cells.***
A. B16 tumor cells show enhanced proliferation in response to CS1. N=8/8, **p<0.01.
B. CS1 inhibits early and late apoptosis in B16 cells as measured using annexinv propidium iodide staining, but the degree of inhibition albeit statistically significant is small. N=20/20. *p<0.05, **p<0.01.
C-D. Stromal cells treated similarly failed to show any increase in proliferation (C) or change in apoptosis (D). N=13/13.

Methods:
Cells were treated with CS1 at a final concentration of 20 µg/ml for 24 hours before evaluation for proliferation and apoptosis. Afterwards the cells were detached using cell dissociation buffer and washed with PBS. To determine proliferation, the cells were

first fixed for 10 minutes in 1% PFA at 4°C and then washed twice. Next, the cells were permeabilized using 0.1% Triton-X-100 for 10 minutes at 4°C followed by 2 additional washing steps. The cells were exposed to PBS containing 5% FCS for 15 minutes at 4°C in order to block non-specific staining. The cells were then washed, stained with an antibody against Ki67 labelled with PE (1:100) for 30 minutes at 4°C, washed again and then analyzed by flow cytometry. To determine apoptosis, the cells were stained against annexin-v labeled with alexa 647 (#640912 Biolegend, 1:50) in annexin-binding-buffer containing 10mM HEPES pH 7.4, 140mM NaCl and 2.5mM $CaCl_2$ for 20 minutes before stopping the reaction with an equal volume of the binding buffer. Five minutes before measurements, propidium iodide was added to the cells.

### Supplementary Figure 5

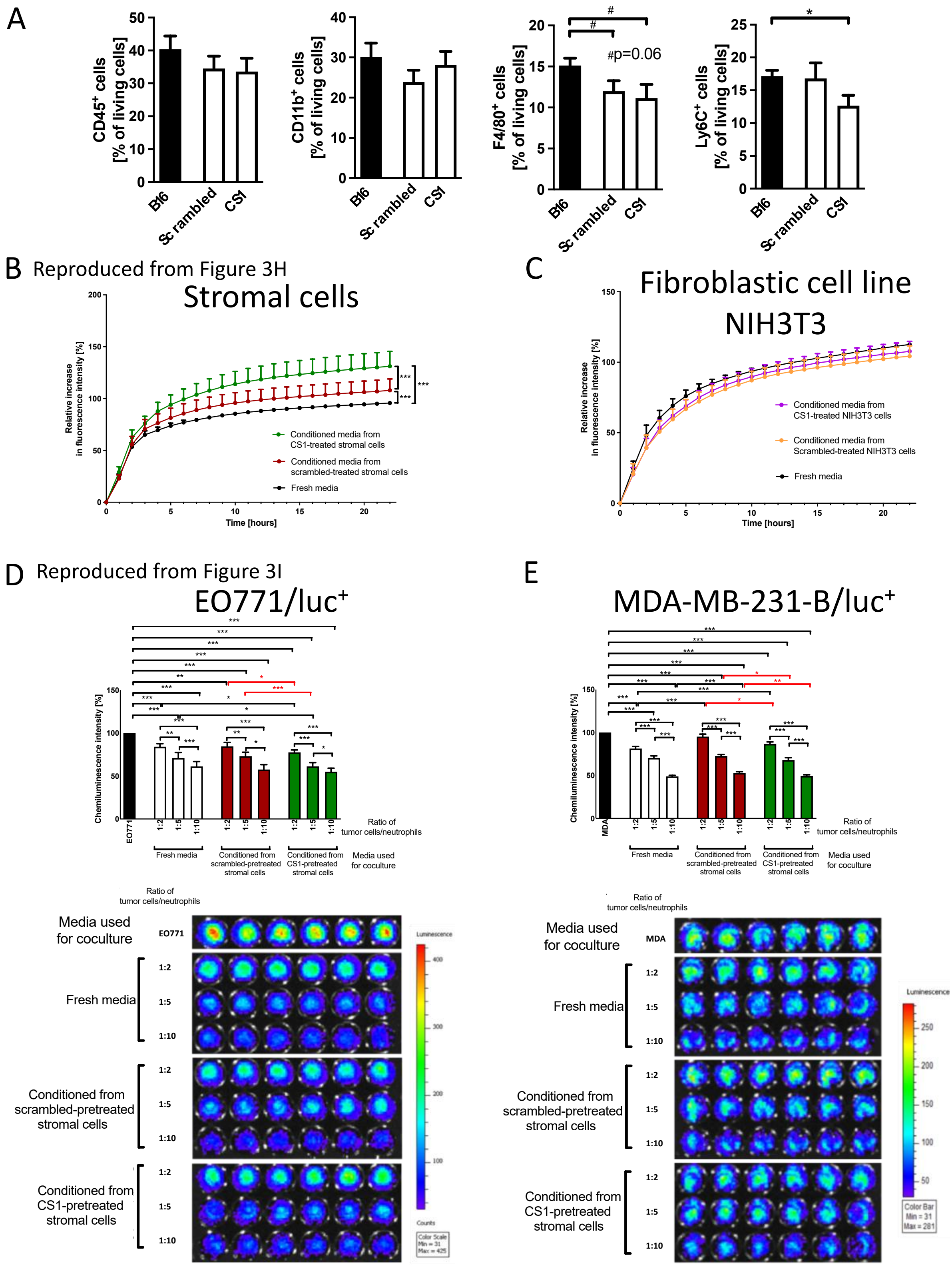


***Supplementary Figure 5.***
***A. Immune cell changes in the presence of stromal cells***

The percentage of various immune cell types was evaluated in tumors exposed to stromal cells that were pretreated with scrambled peptide or CS1 and no statistically significant differences were detected, except in Ly6C-expressing cells. N=12/10/11, *p<0.05.

***B-E. CS1 enhances Ly6G+ cell migration and affects activity in vitro***

B. (Reproduced from Figure 3H for comparison). CS1 enhances migration of Ly6G$^+$ cells towards conditioned media from CS1-pretreated stromal cells (in green) compared to scrambled-pretreated cells (in red). N=11/11/11; ***p<0.001. C. Conditioned media from a fibroblastic cells NIH3T3 fail to cause the same increase. Therefore, increased transmigration of ly6G$^+$ cells towards conditioned media is specific to stromal cells as shown in B. N=5/5/5. D. (reproduced from Figure 3I to allow comparison and show representative data). Cytotoxicity assay using Ly6G$^+$ cells isolated from the bone marrow and cocultured with EO771/luc$^+$ tumor cells in the presence of conditioned media shows statistical differences. The comparisons in red, show that the amount of cancer cells differed between CS1- (green bars) and scrambled-pretreated conditioned media (red bars). This is in line with smaller tumors in CS1-pretreated conditioned media, suggesting higher killing efficacy of Ly6G$^+$ cells, but this effect is small. Note that less remaining tumor cells results in lower luminescence signal intensity. N= 18/18/18/18/18/18/18/18/18/18 , *p<0.05, **p<0.01, ***p<0.001. E. Cytotoxicity assay using Ly6G$^+$ cells isolated from the bone marrow and cocultured with MDA-MB-231-B/luc$^+$ tumor cells in the presence of conditioned media shows statistical differences. The comparisons in red, show the statistical differences between pretreatment with CS1 (in green) and scrambled peptide (in red). These comparisons are consistent with the smaller tumors in the presence of CS1-pretreated cells suggesting higher killing efficacy of Ly6G$^+$ cells, but this effect is small. N= 12/12/12/12/12/12/12/12/12/12 , *p<0.05, **p<0.01, ***p<0.001. ANOVA, if significant, was followed by t-test comparisons.

Methods:

Schematic explaining the experiment was shown in Figure 3G. The design of the experiments is as follows: To evaluate migration and cytotoxicity of Ly6G$^+$ cells, freshly isolated stromal cells were pretreated with CS1. 24 hours later, fresh medium was added for another 24 hours. Conditioned media were applied in the lower well in a transwell assay in the migration assay and migration of fluorescently labeled neutrophils through the floor of the insert was evaluated by measuring the fluorescence in the lower well. Conditioned media were also added to the mixture of luciferase-expressing tumor cells and freshly isolated Ly6G$^+$ cells at different ratios. 24 hours later, the luminescence signal of the remaining tumor cells was evaluated by adding luciferin.

#### Supplementary Figure 6

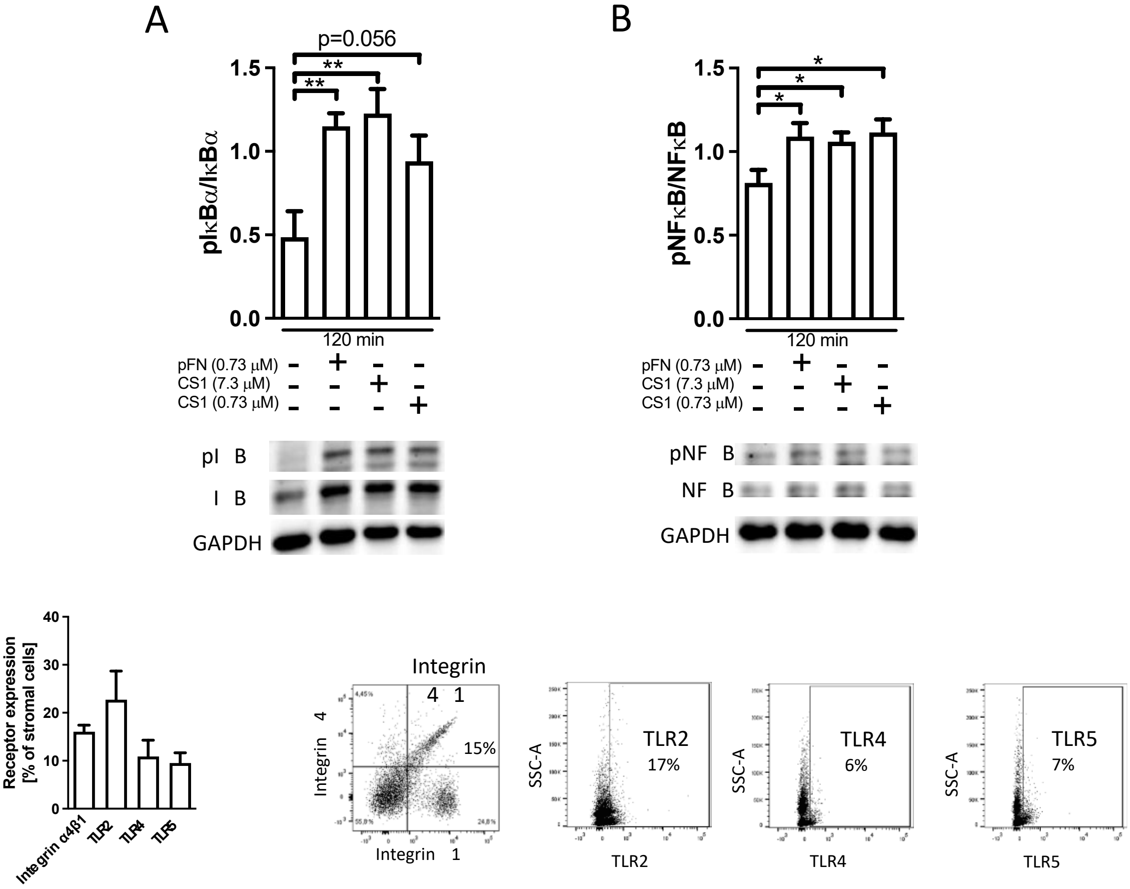


***Supplementary Figure 6.***
***A-B. CS1 increases NF-κB signaling.***
A. Phosphorylation of IκB in response to pFN or CS1 addition to stromal cells was increased, but did not reach statistical significance for the CS1 concentration that was equivalent to the pFN concentration. ANOVA was performed and followed by t-tests. N=8/8/8/8; $^{**}p<0.01$. B. Phosphorylation of NF-κB in response to pFN and CS1 addition to stromal cells is increased with treatment. ANOVA was performed and followed by t-tests. N=8/8/8/8 , $^{*}p<0.05$. Freshly isolated stromal cells were starved overnight, exposed to 160 µg/ml plasma fibronectin and 20 µg/ml and 2 µg/ml of CS1 for 2 hours and nuclear extracts were generated and evaluated by western blotting.

***C. Expression of some of the receptors known to mediate fibronectin effects on cells and to interact with NF-κB.*** TLRs are expressed on up to roughly 20% of the stromal cells, while integrin $\alpha 4\beta 1$ is expressed on less cells. N=3 replicates. Stromal cells were evaluated by flow cytometry.

***Supplementary Table 1. List of differentially expressed molecules between osterix expressing cells and those not expressing osterix***

Only molecules with p<0.01 and ratio difference of > 2 are shown. N= 4/4. Cells were isolated from the bone marrow of mice with the genotype osterix-cre_tdTomato$^{fl/+}$. Based on expression of osterix-cre, tomato-stained cells were detected by flow cytometry and sorted. These were compared to those that did not express tomato using proteomic analysis. The table differs from the volcano plot, because in the table, cells that had no values did not lead to discarding the molecule, as long as a t-test could be performed.

| Protein names | Gene names | Mean CT | Mean Osx | Ratio Osx/CT | p-value | N |
|---|---|---|---|---|---|---|
| Prolyl endopeptidase FAP;Antiplasmin-cleaving enzyme FAP. soluble form | Fap | 2120 | 12572 | 5.93 | 0.0008 | 3/3 |
| Angiopoietin-related protein 3 | Angptl3 | 5202 | 27025 | 5.20 | 0.0004 | 4/4 |
| Contactin-1 | Cntn1 | 11153 | 25327 | 2.27 | 0.0019 | 4/4 |
| Interleukin-1 receptor accessory protein | Il1rap | 24149 | 51454 | 2.13 | 0.0083 | 4/4 |
| Fumarylacetoacetase | Fah | 5054 | 10693 | 2.12 | 0.0069 | 3/3 |
| TIR domain-containing adapter molecule 2 | Ticam2 | 622628 | 1228975 | 1.97 | 0.0028 | 4/4 |
| Collagen alpha-1(I) chain | Col1a1 | 288188 | 558155 | 1.94 | 0.0014 | 4/4 |
| Flavin reductase (NADPH) | Blvrb | 154853 | 295400 | 1.91 | 0.0025 | 4/4 |
| RB binding protein 7, chromatin remodeling factor | Rbbp7 | 105357 | 192510 | 1.83 | 0.0094 | 4/4 |
| PWWP domain-containing protein MUM1L1 | Mum1l1 | 371378 | 665340 | 1.79 | 0.0030 | 4/3 |
| Immunity-related GTPase family Q protein | Irgq | 11503675 | 19140750 | 1.66 | 0.0098 | 4/4 |
| Signal recognition particle 14 kDa protein;Signal recognition particle 14 kDa protein. N-terminally processed | Srp14 | 4375 | 2754 | 0.63 | 0.0026 | 2/3 |
| Ras-related protein Rab-8A | Rab8a | 13735 | 6537 | 0.48 | 0.0004 | 3/2 |
| 26S protease regulatory subunit 6A | Psmc3 | 13807 | 4688 | 0.34 | 0.0003 | 4/2 |
| Heat shock protein HSP 90-alpha | Hsp90aa1 | 104322 | 33588 | 0.32 | 0.0071 | 4/4 |
| Hsc70-interacting protein | St13 | 25308 | 7642 | 0.30 | 0.0029 | 4/4 |
| Nuclear migration protein nudC | Nudc | 15657 | 4612 | 0.29 | 0.0007 | 3/3 |
| Phenylalanine--tRNA ligase alpha subunit | Farsa | 13714 | 2811 | 0.20 | 0.0080 | 3/2 |
| Nucleoplasmin-3 | Npm3 | 20503 | 4011 | 0.20 | 0.0081 | 3/2 |

| | | | | | | |
|---|---|---|---|---|---|---|
| Ran-specific GTPase-activating protein | Ranbp1 | 30045 | 5215 | 0.17 | 0.0034 | 4/4 |
| Peptidyl-prolyl cis-trans isomerase FKBP4;Peptidyl-prolyl cis-trans isomerase FKBP4. N-terminally processed;Peptidyl-prolyl cis-trans isomerase | Fkbp4 | 23238 | 3995 | 0.17 | 0.0023 | 3/3 |
| T-complex protein 1 subunit zeta | Cct6a | 32951 | 5473 | 0.17 | 0.0031 | 3/2 |
| Hypoxanthine-guanine phosphoribosyltransferase | Hprt1 | 27119 | 3881 | 0.14 | 0.0065 | 4/4 |
| ELAV-like protein 1 | Elavl1 | 43844 | 5320 | 0.12 | 0.0007 | 3/3 |
| Small ubiquitin-related modifier;Small ubiquitin-related modifier 2;Small ubiquitin-related modifier 3 | Sumo3; Sumo2 | 67595 | 7391 | 0.11 | 0.0015 | 3/2 |
| Thymosin beta-10 | Tmsb10 | 178900 | 17933 | 0.10 | 0.0091 | 3/3 |
| T-complex protein 1 subunit eta | Cct7 | 36197 | 2537 | 0.07 | 0.0000 | 3/2 |
| ATP-dependent RNA helicase A | Dhx9 | 30496 | 1839 | 0.06 | 0.0014 | 3/2 |
| Multifunctional protein ADE2;Phosphoribosylaminoimidazole-succinocarboxamide synthase;Phosphoribosylaminoimidazole carboxylase | Paics | 16929 | 411 | 0.02 | 0.0041 | 3/2 |
